# Mapping the Genetic Landscape of DNA Double-strand Break Repair

**DOI:** 10.1101/2021.06.14.448344

**Authors:** Jeffrey A. Hussmann, Jia Ling, Purnima Ravisankar, Jun Yan, Ann Cirincione, Albert Xu, Danny Simpson, Dian Yang, Anne Bothmer, Cecilia Cotta-Ramusino, Jonathan S. Weissman, Britt Adamson

## Abstract

Cells repair DNA double-strand breaks (DSBs) through a complex set of pathways that are critical for maintaining genomic integrity. Here we present Repair-seq, a high-throughput screening approach that measures the effects of thousands of genetic perturbations on the distribution of mutations introduced at targeted DNA lesions. Using Repair-seq, we profiled DSB repair outcomes induced by two programmable nucleases (Cas9 and Cas12a) after knockdown of 476 genes involved in DSB repair or associated processes in the presence or absence of oligonucleotides for homology-directed repair (HDR). The resulting data enabled principled, data-driven inference of DSB end joining and HDR pathways and demonstrated that repair outcomes with superficially similar sequence architectures can have markedly different genetic dependencies. Systematic interrogation of these dependencies then uncovered unexpected relationships among DSB repair genes and isolated incompletely characterized repair mechanisms. This work provides a foundation for understanding the complex pathways of DSB repair and for optimizing genome editing across modalities.

## INTRODUCTION

DNA double-strand breaks (DSBs)—lesions that sever the phosphate backbones of two complementary DNA strands—pose a serious threat to cells and organisms. Failure to repair DSBs, which arise from either exogenous or endogenous sources, can have deleterious consequences, including genome instability and cell death^1^. To protect themselves, cells have evolved a sophisticated, multifaceted system of DSB repair with many iterative and redundant mechanisms, each capable of contributing to repair, albeit with different tendencies towards a variety of sequence outcomes (e.g., error-free repair, insertions, deletions). Given that defects in these mechanisms underlie human diseases, including cancer, there has long been interest in characterizing them; however, our understanding of the complex landscape of DSB repair mechanisms remains far from complete.

To synthesize our knowledge of DSB repair into a single framework, repair mechanisms can be conceptualized as a decision tree, with “branches” representing the various molecular events leading to a repaired break and “nodes” indicating points of transition or commitment between processes^2, 3^. In accordance with this characterization, there are two major pathways of DSB repair, which are separated by clear regulatory mechanisms. The first of these pathways, non-homologous end joining (NHEJ), comprises a set of flexible end-joining mechanisms that act throughout the cell cycle to facilitate the processing, synapsis, and direct ligation of DSB ends. The second, homology-directed repair (HDR), uses homologous templates to facilitate repair in S/G2 after 5’-to-3’ nucleolytic processing of DSB ends. Despite the utility of this conceptual framework, however, even the most carefully constructed models of DSB repair remain incomplete, with gaps and ambiguities in pathway organization. The events upstream and downstream of the bifurcation between NHEJ and HDR, for example, remain incompletely defined, obscuring the total number of distinct routes to repair. Moreover, alternative branches of DSB repair, loosely described as alternative end joining, have been difficult to fully characterize. Indeed, it remains unclear whether these mutagenic processes represent one or multiple overlapping pathways in human cells^4^. In this regard, existing approaches for studying DSB repair are limiting, because they rely primarily on phenotypes that read out only single repair outcomes (i.e., reporters)^5, 6^, indirectly measure the activities of many pathways (i.e., sensitivity to genotoxic agents), or can examine only a few perturbations at a time (i.e., targeted sequencing)^4, 7–9^. Yet, any effort to comprehensively map DSB repair pathways should ideally both measure the full spectrum of repair outcomes and define their genetic dependencies.

To address this shortcoming, we developed Repair-seq, a scalable and quantitative method for systematically delineating mechanisms of DNA repair. This platform combines locus-specific deep sequencing with CRISPR-interference (CRISPRi)-based genetic screens to measure the effects of thousands of genetic perturbations on the spectrum of DNA repair outcomes produced by the introduction of a targeted DNA lesion. Paired with an analytical pipeline for characterizing and quantifying repair outcomes, Repair-seq generates high-information-content readouts of DNA repair for many genetic regulators simultaneously, allowing both systematic classification of repair outcomes and delineation of gene functions. Using Repair-seq, we then constructed principled, data-driven maps of DSB repair outcomes induced by Cas9 and Cas12a and their genetic dependencies. Systematic exploration of these maps nominated novel repair mechanisms and revealed new roles for canonical DNA damage response genes. As examples, we identified a role for *RAD17* and genes encoding components of the 9-1-1 complex in specifically promoting *POLQ*-mediated repair outcomes, and we delineated several distinct mechanisms that generate insertions during repair of Cas9-induced DSBs.

Beyond shedding light on the diversity of mechanisms used by cells to repair DSBs, our efforts demonstrate that Repair-seq can be used to study genome editing technologies, most of which rely on DNA damage to install sequence changes. As a proof of principle, we used Repair-seq to study homology directed repair with single-stranded oligonucleotide donors. This effort revealed a range of mechanisms by which donor sequence can be incorporated at breaks in both intended and unintended ways. Altogether, these data and associated insights provide a foundation for further exploration of DSB repair mechanisms and demonstrate the broad utility of Repair-seq for investigating the repair processes engaged by different genome editing technologies.

## RESULTS

### A functional genomics platform for interrogating DSB repair pathways

Enzymatically induced DSBs generate diverse but reproducible sets of mutations, typically in the form of short insertions and deletions (indels)^9–14^. Targeted sequencing of these mutations after genetic or chemical perturbations have revealed coordinated shifts in their frequencies^4, 7–9^, indicative of the fact that each repair outcome is produced by a specific set of processing events and therefore promoted or suppressed by the activity of specific gene products. Genetic screens have been useful for identifying these genes, but assays amenable to high-throughput screening typically only measure repair products associated with single repair pathways (e.g., reporter assays^15–20)^ or report the combined activity of many DSB repair processes (e.g., growth assays^21–23)^. On the other hand, targeted sequencing of DSB-induced mutations has been limited to low numbers of perturbations.

We reasoned that a screening platform capable of measuring the effects of many genes on each of many DSB-induced repair products would enable systematic interrogation of DSB repair mechanisms. We built this platform, called Repair-seq, by combining high-throughput CRISPR-based genetic screening with targeted sequencing of DSB repair outcomes (Figure 1A and 1B). CRISPR-based screens use libraries of single-guide RNAs (sgRNAs) to enable massively parallel interrogation of gene function in scalable, pooled experiments^24–26^. To take advantage of this scalability, we designed a screening vector that links an sgRNA expression cassette to a nearby target region for DNA damage (Figures 1A and S1A). After genomic integration, induction of damage within this region pairs the products of DNA repair to perturbation identifiers (i.e., sgRNA sequences). This association, in turn, enables linked recovery and paired-end sequencing of those features and thus measurement of the frequency of repair outcomes within populations of cells that received each perturbation (Figure 1B and Method Details).

**Figure 1.**
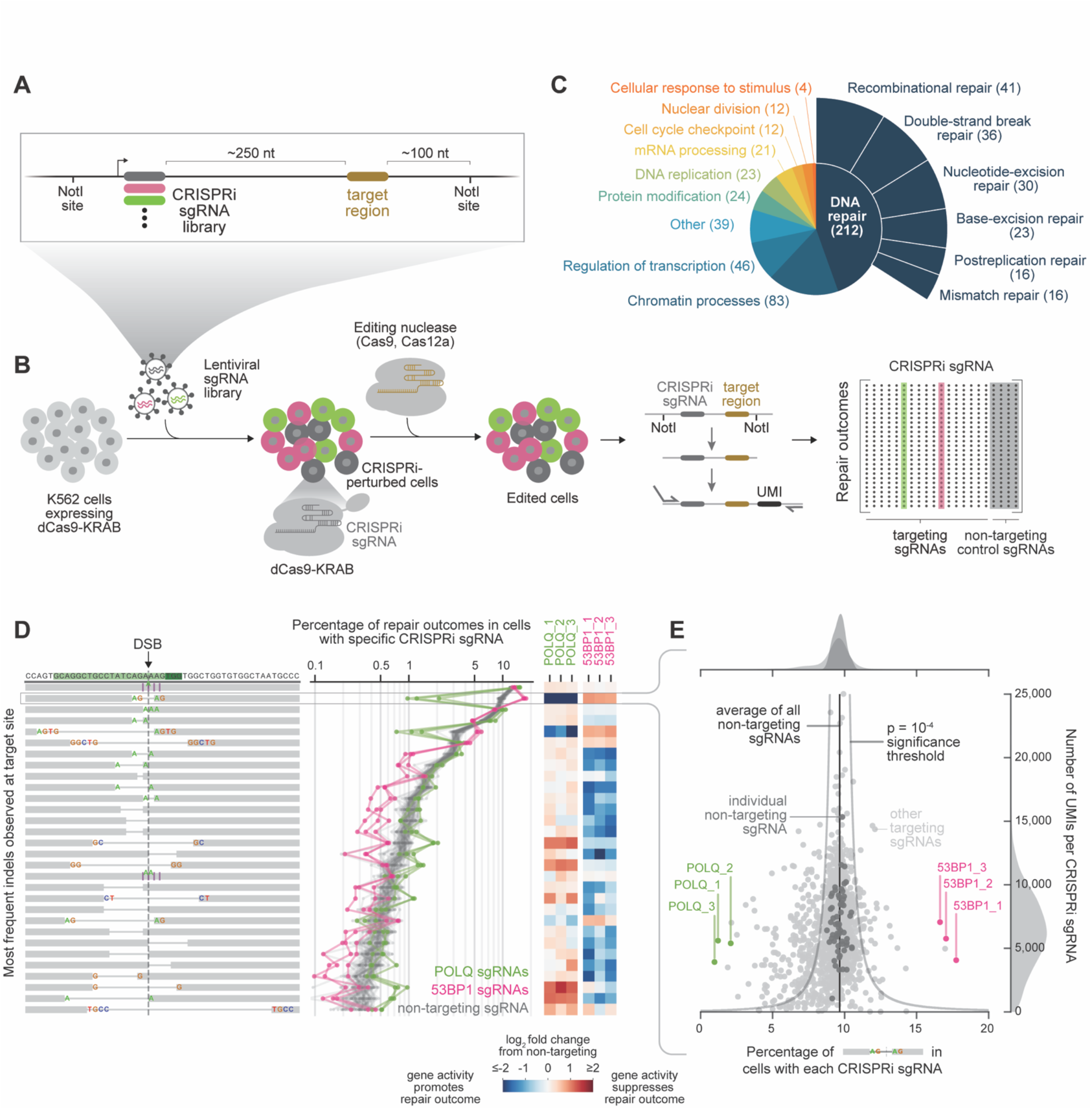
Repair-seq is a high-resolution screening platform for systematically interrogating DNA repair processes. (A) Schematic of screening vector with linked CRISPRi sgRNA expression cassette and target region for induced DNA damage. (B) Experimental workflow of a Repair-seq screen. Cells expressing a CRISPRi effector protein (dCas9- KRAB) are infected with CRISPRi sgRNAs linked to a targetable editing region. After allowing time for CRISPRi to repress targeted gene expression, a programmable nuclease directed at the editing site is introduced into the pool of perturbed cells by electroporation. The genomic region containing the CRISPRi sgRNA and repair outcome is later isolated, ligated with a unique molecular identifier (UMI), and amplified. Paired-end sequencing of linked CRISPRi sgRNA identities and repair outcomes is performed to measure perturbation-specific repair outcome distributions. (C) Functional annotation classes for genes targeted by 1,573 sgRNA CRISPRi library (see also Table S5). (D) Data from initial Repair-seq screen viewed from a CRISPRi-sgRNA-centric perspective. Diagrams (left) show the 30 most frequent indels observed in cells receiving non-targeting CRISPRi sgRNAs. Green rectangles in the top row mark the protospacer and PAM of Cas9 target site. Vertical dashed line marks the expected DSB location. Purple vertical lines with colored sequence indicate insertions. Grey horizontal lines indicate deletions, with any flanking microhomology indicated by colored sequence. Plot (middle) shows frequency of each outcome in the presence of individual non-targeting sgRNAs (60 grey lines) or sgRNAs targeting *POLQ* (3 green lines) or *53BP1* (3 pink lines). Heatmaps (right) display log_2_ fold changes in the frequency of each outcome for *POLQ* or *53BP1* sgRNAs relative to average frequency across all non-targeting sgRNAs. (E) Data from initial Repair-seq screen viewed from an outcome-centric perspective, focusing on one microhomology-flanked 4 nt deletion. Plot shows number of UMIs recovered for each sgRNA (y-axis) against the percentage of UMIs reporting this specific deletion (x-axis). Green dots mark sgRNAs targeting *POLQ*, pink dots mark sgRNAs targeting *53BP1*, dark grey dots mark individual non-targeting sgRNAs, and light grey dots mark all other sgRNAs. Black vertical line indicates the average percentage across all non-targeting sgRNAs. Dark grey density plot above the scatter plot shows a kernel density estimate (KDE) of the distribution of frequencies in non-targeting sgRNAs; light grey density plots above and to the right show KDEs of the distribution of frequencies (top) and UMI counts (right) for all sgRNAs. The indicated significance threshold marks the deviation in frequency expected to occur by chance with probability less than 10^-4^ for each number of UMIs given the observed average non-targeting frequency under a binomial model.

We tested induction and repair of DSBs within the context of our screening construct using a target region derived from the endogenous *HBB* locus (Figure S1A). After transducing this vector into the human myelogenous leukemia cell line K562, we induced DSBs at several locations (or “target sites”) within the target region by electroporation of different *Streptococcus pyogenes* Cas9 RNPs (Figure S1A and Table S1). After 3 days of recovery, we sequenced repair junctions for each target site in the integrated construct and at the endogenous *HBB* locus, revealing diverse distributions of DSB repair outcomes (Figure S1B and S1C). These distributions were highly reproducible within both the integrated construct and the endogenous locus but differed considerably between distinct target sites, including sites located only 1 base pair (bp) apart (Figure S1A and S1D). Critically, despite the potential influence of local chromatin context^27^, distributions from the integrated *HBB* target region were closely concordant with those from the endogenous *HBB* (Figure S1E). We therefore concluded that repair outcomes generated within our integrated screening construct would produce reproducible and highly-informative readouts of DSB repair.

In developing our assay, we also considered how to accurately quantify the effects of genetic perturbations on a broad range of repair outcomes. Because many genes with known roles in DNA repair and replication are essential, we chose to perturb gene expression with CRISPRi (Figure 1B), a system of gene inactivation that uses a catalytically-inactive Cas9 fusion protein (dCas9-KRAB) to inhibit transcription at targeted promoters^28, 29^. CRISPRi homogeneously represses expression of targeted genes across cell populations^29^ and has previously been shown to be well-suited for investigating genes whose inhibition causes adverse growth phenotypes^24^. Next, to ensure accurate measurement of repair outcomes, we developed a strategy for preparing sequencing libraries in which unique molecular identifiers (UMIs) are attached to repair outcomes prior to amplification (Figures 1B and S1F). These UMIs allow us to collapse reads generated from the same repair product, effectively turning the assay into a single-molecule measurement in which repair events from each cell are counted only once^30^. This process also allows for correction of errors introduced by PCR or sequencing^31^, and thus enables us to distinguish infrequent but real repair outcomes from technical artifacts (Method Details). Finally, to maximize the scope of repair outcomes analyzed, we developed a computational outcome classification pipeline based on our previously described *knock-knock* approach^32^. This pipeline comprehensively characterizes repair outcomes, including indel architectures and outcomes consisting of more complex sequence rearrangements (Method Details). We note however that repair outcomes without priming regions necessary for sequencing library construction (e.g. sufficiently large deletions^33^) would not be detected (Figure S2A-S2E and Method Details).

### Repair-seq enables sequence-independent classification of DSB repair outcomes and systematic identification of functionally related genes

Many previous efforts have identified genes involved in DNA repair in human cells^16, 18–21, 34–36^. Taking advantage of this knowledge, we designed a custom sgRNA library targeting a set of 476 genes with roles in DNA repair or associated processes (Figure 1C and Table S2). This library of 1,573 total CRISPRi sgRNAs includes typically 3 sgRNAs targeting each gene and 60 non-targeting control sgRNAs. After cloning this library into the *HBB*-derived screening vector, we performed an initial Repair-seq screen in K562 cells expressing dCas9-KRAB (Table S3 and Method Details). During screening, we allowed 6 days for CRISPRi-mediated gene depletion, then electroporated cells with one *HBB*-targeting Cas9 RNP and allowed 3 days for induction and repair of DSBs. We then extracted genomic DNA and prepared sequencing libraries to measure the distribution of repair outcomes for each genetic perturbation.

To quantify phenotypes from our screen, we compared the frequency of each DSB repair outcome in the presence of a gene-targeting CRISPRi sgRNA to the average frequency of the same outcome across all non-targeting control sgRNAs (Figures 1D and S2F). A reduction in the frequency of an outcome upon knockdown of a gene is evidence that activity of the gene’s product promotes formation of the outcome, while an increase indicates that the gene’s product suppresses the outcome. Enabling these comparisons, frequencies of individual DSB repair outcomes were highly reproducible between all 60 non-targeting controls in our library (represented by gray lines in Figures 1D and S2F). Averaged together, these measurements produced a high-confidence baseline outcome distribution in unperturbed cells. Moreover, performing the screen at extremely high coverage (median 6,312 UMIs recovered per sgRNA; Table S3) enabled statistically significant comparisons across many repair outcomes, including those produced at low frequency (Figures 1D, 1E, and S2G).

We explored data from this initial Repair-seq screen from two complementary perspectives. First, we examined the effects of many different gene knockdowns on specific repair outcomes (Figures 1E and S2G), producing parts lists of genes involved in creating or preventing each outcome. For example, a high-frequency deletion with two nts of microhomology (MH) at the junction site was depleted by sgRNAs targeting *POLQ*, a gene known to promote deletions mediated by paired microhomologies^37, 38^, and increased by sgRNAs targeting *53BP1*, a gene that represses *POLQ*-mediated end joining^39^ (Figure 1E). Second, we examined the effects of specific CRISPRi sgRNAs across many different repair outcomes (Figures 1D and S2F), producing detailed views of the role that each targeted gene plays in different repair pathways. Viewed from this perspective, the effect of *POLQ* knockdown across all deletions with microhomology was unexpectedly complex, with knockdown depleting a specific subset of these deletions but increasing or having no effect on others (Figure 1D).

In another example, CRISPRi sgRNAs targeting *XRCC5* and *XRCC6,* genes that encode components of a heterodimer with central roles in NHEJ^40^, produced specific increases in two different short insertions (Figure S2F). This example demonstrated that loss of gene products with functionally similar roles can produce strikingly similar redistributions of outcomes, suggesting that systematic comparison of sgRNA outcome redistribution signatures would allow identification of other functionally related genes. Conversely, the observation that outcomes with related sequence characteristics sometimes shared genetic regulators (e.g., the subset of MH-flanked deletions promoted by *POLQ* and suppressed by *53BP1* in Figure 1D, and the two insertions suppressed by *XRCC5* and *XRCC6* in Figure S2F) suggested that by comparing the genetic dependencies of different repair outcomes, we could identify repair products produced by similar mechanisms. We therefore reasoned that hierarchical clustering of Repair-seq phenotypes along both dimensions could produce a high-resolution map of DSB repair processes.

To test this idea, we first focused on a subset of our data, using repair outcomes produced above a baseline frequency of 0.2% (retaining 48 outcomes) and the 100 sgRNAs that caused the strongest redistribution of these outcomes (Method Details). We formed an outcome-by-sgRNA matrix of the log_2_ fold changes in outcome frequencies caused by each sgRNA relative to the average of all non-targeting sgRNAs and clustered along both dimensions (Figure 2). Clustering of sgRNAs revealed a rich set of distinct signatures across our library, with sgRNAs targeting the same gene grouped closely together and sgRNAs against genes with known functional relationships also often placed together (e.g. *XRCC5* and *XRCC6*; *BARD1* and *BRCA1*). Clustering of repair outcomes, on the other hand, revealed groups of outcomes with closely related genetic dependencies, especially among those with similar sequence features (e.g., insertions, deletions of sequence on only one or both sides of the break). Below, we expand this analysis, demonstrate the validity and reproducibility of the approach, and discuss insights gained from its application.

**Figure 2.**
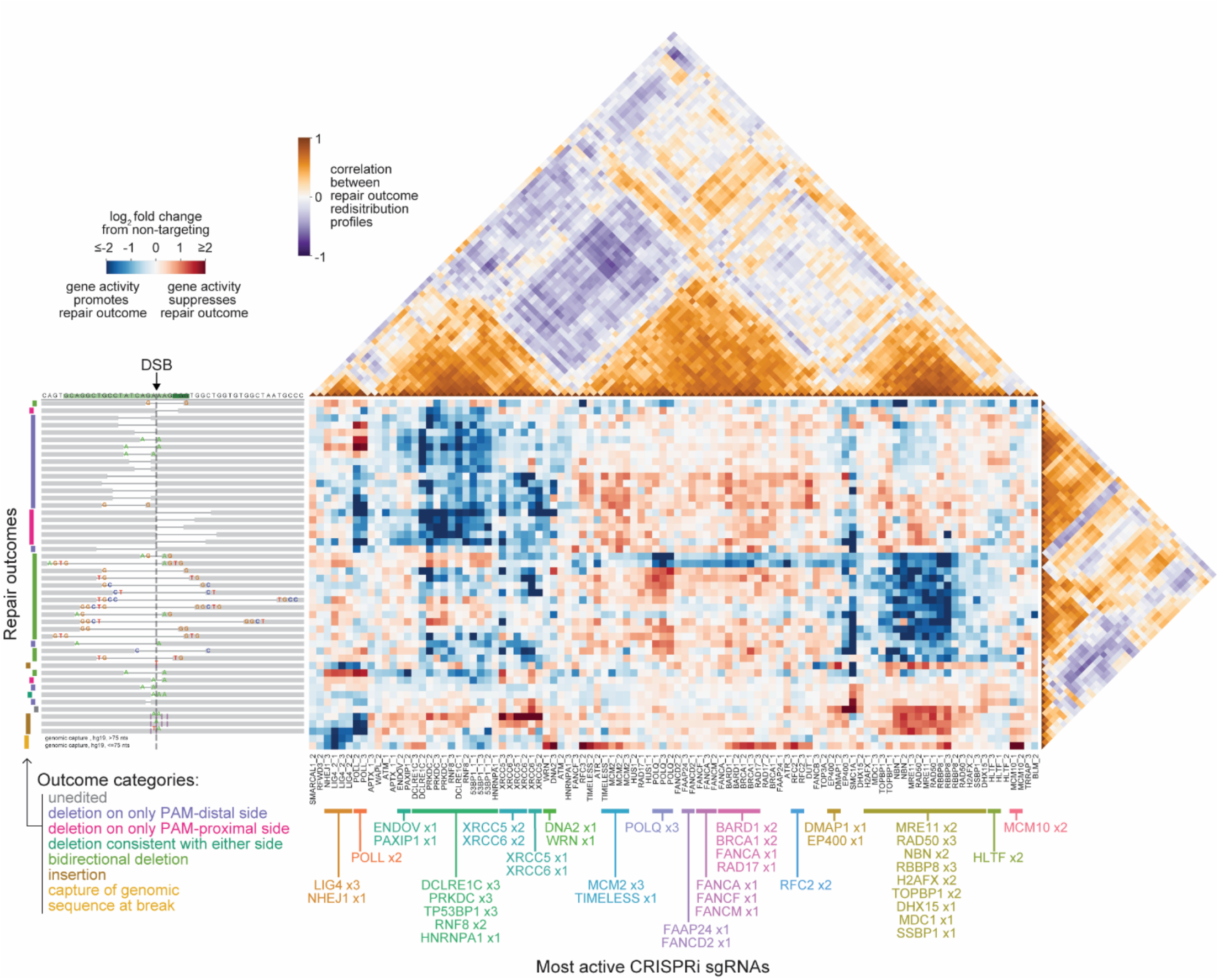
Repair-seq enables data-driven inference of the genetic organization of DSB repair mechanisms. Central heatmap displays log_2_ fold changes in the frequencies of repair outcomes for the 100 most active CRISPRi sgRNAs relative to the average of all non-targeting sgRNAs in initial Repair-seq screen, hierarchically clustered along both dimensions. Rows of the central heatmap correspond to distinct outcomes (i.e., all individual sequence outcomes present above baseline frequency of 0.2% and two composite outcomes representing the combined frequency of all outcomes in which stretches of genomic sequence <75 nts or >75 nts were captured at the break). Outcomes are depicted by diagrams on the left. Columns of the central heatmap correspond to distinct CRISPRi sgRNAs. Triangular heatmaps depict correlations between pairs of sgRNAs (above) and between pairs of outcomes (right). sgRNA cluster assignments produced by HDBSCAN are labeled below.

### Repair-seq screens at multiple target sites produce a high-resolution map of DSB repair

Data from previous reports^9–14^, as well as our own data (Figure S1D), have indicated that local sequence context is a strong determinant of DSB repair outcomes. Repair of DSBs within any particular sequence may therefore provide an incomplete or idiosyncratic view of the full range of possible repair mechanisms. To address this possibility, we applied Repair-seq to a broader set of Cas9-induced repair outcomes by screening DSBs at additional locations within our integrated screening construct (Figures 3A and S3A).

**Figure 3.**
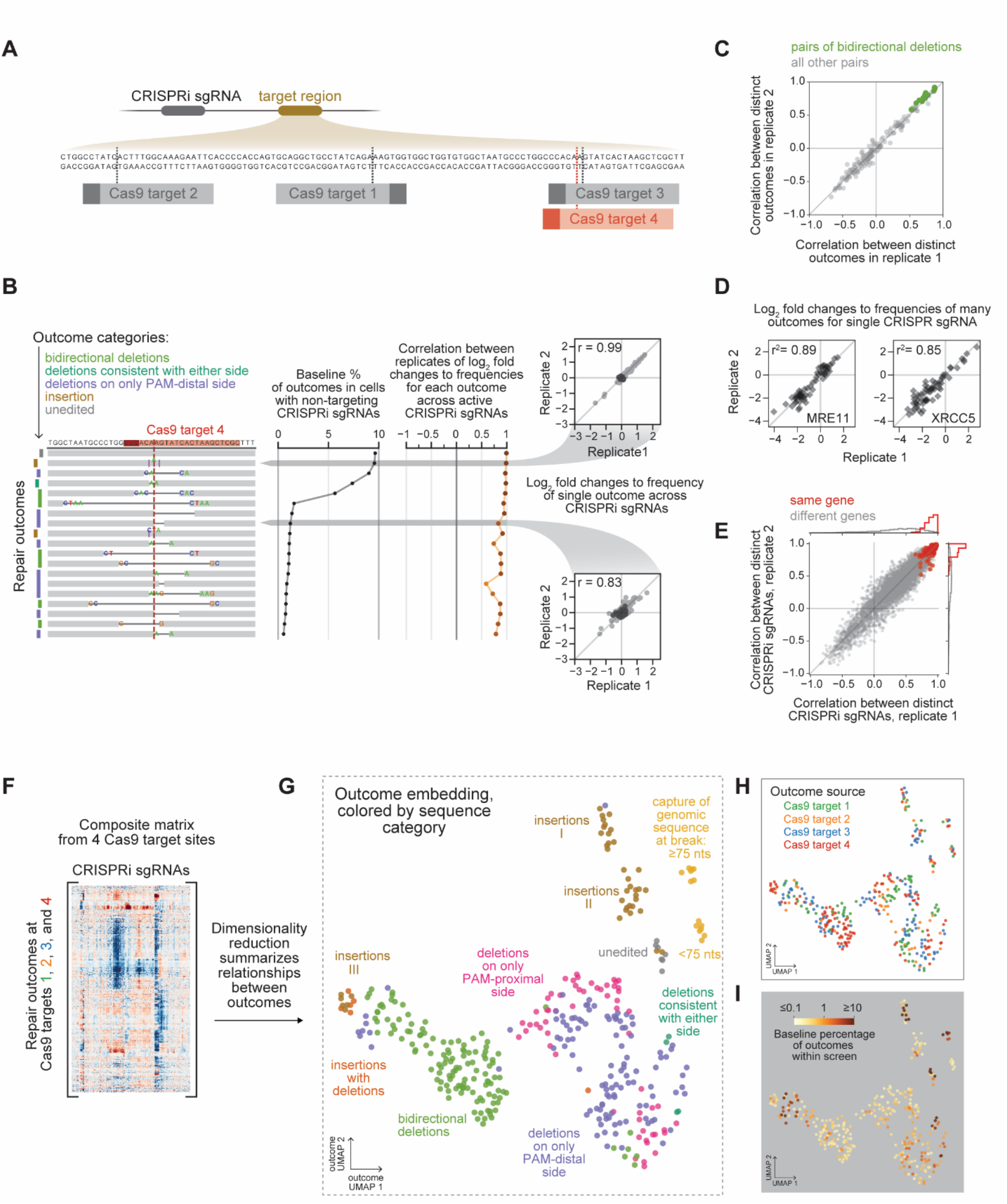
A systematic map of the genetic dependencies of repair outcomes generated at Cas9- induced DSBs. (A) Cas9 target sites in the Repair-seq screening construct. Data from screens at target site 4 (red) are shown in panels B-E. Data from screens at other target sites are shown in Figure S3A and S3E-S3M. (B) Evaluation of reproducibility of the effects of CRISPRi sgRNAs on individual outcome frequencies. Diagrams (left) depict most frequent repair outcomes observed for Cas9 target site 4. Graphs show baseline percentages of each outcome across all non-targeting sgRNAs, averaged across two replicates (middle), and correlation between replicates in log_2_ fold changes to frequencies of corresponding outcome across all sgRNAs that produce a significant overall redistribution of outcomes (right). Insets show comparisons between two replicates of log_2_ fold changes in frequency of one common insertion (top) or less common deletion (bottom) for all CRISPRi sgRNAs relative to the average of non-targeting sgRNAs. Dark grey points mark individual non-targeting sgRNAs. Light grey points mark all other targeting sgRNAs. (C) Correlations between outcome signatures (log_2_ fold changes in frequency of an outcome across active sgRNAs) for pairs of distinct outcomes in replicate 1 (x-axis) and replicate 2 (y-axis). Green points mark pairs of distinct bidirectional deletions. Light grey points mark all other outcome pairs. (D) Comparison of CRISPRi sgRNA signatures (log_2_ fold changes in the frequency of each outcome present above baseline frequency of 0.5% produced by a CRISPRi sgRNA) between two replicates for an sgRNA targeting *MRE11* (left) and an sgRNA targeting *XRCC5* (right). (E) Correlations between CRISPRi sgRNA signatures for distinct sgRNAs in replicate 1 (x-axis) and replicate 2 (y-axis) screens. Red points mark pairs of sgRNAs targeting the same gene. Grey points mark pairs of sgRNAs targeting distinct genes. (F) Representation of composite matrix of log_2_ fold changes in outcome frequencies produced by CRISPRi sgRNAs in screens performed at four different Cas9 target sites (two replicates each for target sites 1, 3, and 4 and one replicate for target site 2). Larger version in Figure S4. (G) UMAP embedding of outcomes from screens at four Cas9 target sites. Each point represents a single outcome in a single screen replicate. Points are colored by outcome sequence architecture category. (H+I) UMAP embedding of outcomes as in G, colored by Cas9 target site of origin (H) or by baseline frequency of outcome within screen of origin (I).

To enable this effort, we built and validated a condensed CRISPRi sub-library consisting of 336 sgRNAs targeting 118 genes and 30 non-targeting control sgRNAs, prioritizing sgRNAs empirically determined to produce significant redistribution of outcome frequencies in initial analyses (Table S4, Figures S3B-S3D, and Method Details). This compact sub-library allowed us to more easily parallelize screens, and using a combination of this sub-library and our 1,573 sgRNA library, we performed screens at four different target sites located across a 102 nucleotide (nt) window of our screening construct using different Cas9 RNPs (Table S1).

Data from these target sites, which were each screened at least twice (Table S3), allowed rigorous evaluation of the technical quality of screen phenotypes. Outcome signatures (i.e., the effects of many CRISPRi sgRNAs on an individual repair outcome) were highly reproducible between screens performed at the same target site, even for lower frequency outcomes (Figures 3B and S3E-S3G). Furthermore, correlations between signatures for distinct outcomes reproducibly spanned a range from high (indicating many shared genetic dependencies between the outcomes) to intermediate (indicating partially-overlapping dependencies) to negative (indicating production by opposing pathways) (Figures 3C and S3H-S3J). Outcome signature correlations therefore represent a high-resolution measurement of relatedness between outcomes. Supporting this interpretation, pairs of outcomes with similar sequence architectures, such as bidirectional deletions, were highly enriched for strong correlations (Figures 3C and S3H-S3J). Similarly, for CRISPRi sgRNAs that caused substantial outcome redistributions, sgRNA signatures (i.e., the effects of an individual sgRNA across many repair outcomes) were highly reproducible across screens performed at the same target sites (Figure 3D), and pairs of sgRNAs targeting the same gene reproducibly produced highly correlated signatures (Figures 3E and S3K-S2M), indicating that these signatures reflect specific effects of gene knockdown.

Given these results, we combined data from screens performed at all four Cas9 target sites into a single composite matrix of log_2_ fold changes in outcome frequencies, with outcomes from each screen at the same target site included individually (Figures 3F and S4). Hierarchical clustering of these data revealed rich structure in patterns of genes with similar effects and outcomes with shared genetic dependencies. To explore these data, we first focused on repair outcomes. Clustering of outcomes frequently grouped identical outcomes from screens performed at the same target site closely together (Figure S4), indicating that the precise dependencies of each DSB repair outcome are highly specific. Nevertheless, outcomes from different screens were also grouped into larger modules.

**Figure 4.**
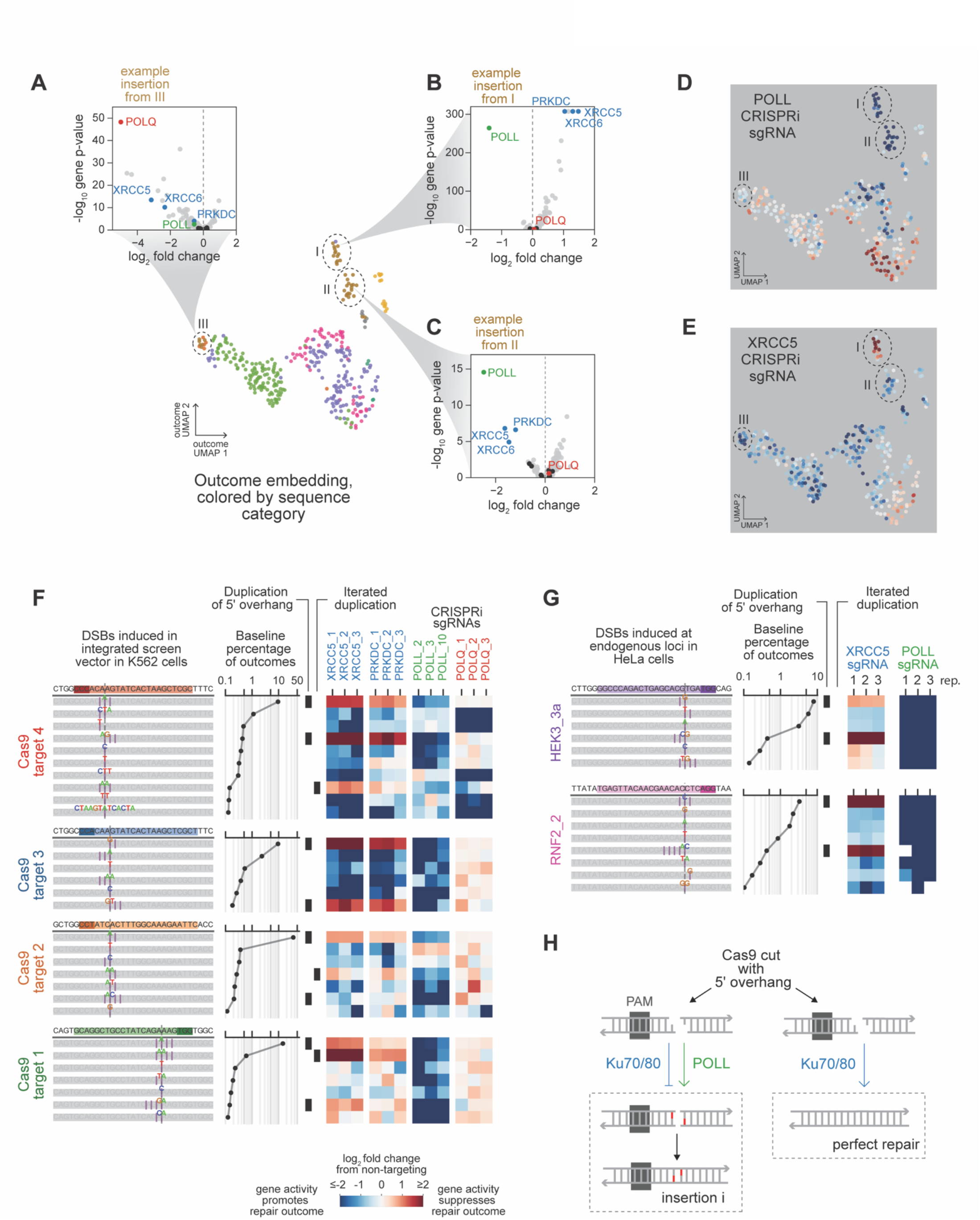
Insertions at Cas9-induced DSBs have distinct sets of dependencies on core NHEJ factors. (A+B+C) Volcano plots depicting gene-level phenotypes of representative insertions from indicated groups in the UMAP embedding of Cas9 outcomes. Phenotypes are calculated as the average log_2_ fold change in frequency produced by the two most extreme sgRNAs per gene. Gene-level p-values are calculated by aggregating binomial p-values from each sgRNA targeting the gene. Black dots represent random sets of three individual non-targeting sgRNAs. (D+E) Log_2_ fold changes in outcome frequencies produced by indicated CRISPRi sgRNAs overlaid on the composite Cas9 UMAP outcome embedding. (F) Effects of *XRCC5-*, *PRKDC-*, *POLL-*, or *POLQ-*targeting sgRNAs on frequencies of insertions at four Cas9 target sites in indicated screens. Diagrams depict all insertions observed at baseline frequency greater than 0.1% at each Cas9 target site (left). Graphs show baseline frequencies of each insertion (middle). Heatmaps display log_2_ fold changes in insertion frequencies produced by indicated sgRNAs. Black bars to the left of heatmaps mark duplications of PAM-distal sequence adjacent to the canonical DSB location (left column) or insertions consistent with multiple iterated duplications (right column). (G) Effects of *XRCC5* and *POLL-*targeting sgRNAs on insertion frequencies at endogenous loci in HeLa cells expressing dCas9-KRAB. Diagrams depict all insertions observed at baseline frequencies greater than 0.1% (left column). Graphs show baseline frequencies of each insertion (middle). Heatmaps display log_2_ fold changes in insertion frequencies produced by indicated sgRNAs in replicates (rep). Black bars to the left of heatmaps mark insertions as in F. (H) Model for generation of Cas9-induced insertions from 5′ overhangs.

To better visualize these modules, we performed dimensionality reduction on the outcome signatures in this aggregate matrix using uniform manifold approximation and projection (UMAP), creating an embedding based on the genetic dependencies of each outcome (Figure 3G). Examination of this DSB repair outcome embedding produced three main observations. First, although outcomes with similar sequence features were often grouped together, many such outcomes were separated into distinct groups, indicative of the fact that outcomes with similar sequence features are often but not always mechanistically related (Figure 3G). For example, small insertions separated into distinct groups (labeled I, II, and III in Figure 3G). A second observation was that outcomes from different screens were interspersed throughout the embedding. This interspersion supports the idea that DSBs induced in different sequences engage broadly similar repair mechanisms (Figure 3H). An interesting exception to this observation was that one group of insertions (III), which was interspersed with more complex insertion+deletion outcomes, arose predominantly from screens performed at one target site (4) (Figures 3G and 3H), suggesting that the underlying mechanism is more sensitive to sequence context (discussed below). Finally, high-frequency outcomes clustered into specific areas of the embedding, highlighting the fact that certain repair processes are preferentially engaged at Cas9-induced breaks^41, 42^, while the genetic dependencies of many individually less frequent outcomes revealed a broader diversity of mechanisms at work (Figure 3I). Altogether, these data represent an in-depth accounting of the mechanisms responsible for Cas9-induced DSB repair.

### Distinct mechanisms for generation of insertions at Cas9-induced DSBs revealed by Repair-seq

A key strength of Repair-seq is the ability to classify repair outcomes by their genetic dependencies rather than assumptions based on their sequences. In our analysis, this strength was highlighted by the separation of short, Cas9-induced insertions into several groups (Figure 3G). Consistent with a general requirement for DNA synthesis in the generation of insertions, insertions exhibited strong dependence on different DNA polymerases, with DNA polymerase theta (Pol θ, encoded by *POLQ*) promoting insertions in one group (III) (Figure 4A) and DNA polymerase lambda (Pol λ, encoded by *POLL*) promoting insertions within the other two (I and II) (Figure 4B-4C). Notably, dependence on *POLL* agrees with a previous study in yeast, in which the Pol λ homolog, DNA polymerase 4 (Pol4), was found to be responsible for Cas9-induced insertions^42^, while the existence of *POLL*-independent insertions (group III, discussed below) highlights that the genetic landscape of insertion mechanisms in human cells is more complex.

Interestingly, bifurcation of *POLL-*dependent insertions into two groups (I and II) suggested that Cas9- induced DSBs engage distinct mechanisms of *POLL-*mediated repair in human cells (Figure 4D). To identify distinguishing genetic determinants, we compared the effects of all gene knockdowns on insertions from each group. This analysis revealed that knockdown of three mechanistically related NHEJ genes, *XRCC5*, *XRCC6,* and *PRKDC,* produced dramatic increases in the frequency of insertions from one of the two *POLL*-dependent groups (I) while decreasing the frequency of insertions from the other such group (II) (Figure 4B-4E). Further examination of insertions produced at the four Cas9 target sites identified common sequence characteristics shared by *XRCC5-*, *XRCC6-*, and *PRKDC-*suppressed insertions. Specifically, all such insertions consisted of one of three sequence types: 1 nt insertions that duplicated the nucleotide on the side of the canonical Cas9 cut site furthest from the protospacer adjacent motif (PAM-distal); 2 nt insertions that duplicated this same PAM-distal nucleotide twice (“iterated duplications”); and 2 nt insertions that duplicated the two nucleotides adjacent to the cut site on the PAM-distal side (Figure 4F). Two of these insertion types are consistent with products of staggered Cas9 cleavage^41–43^. Indeed, although canonically blunt, Cas9-induced DSBs are variably generated with short 5’ overhangs^41, 43^, and fill-in synthesis can create insertions of the overhang sequence^42^. Our results suggest that these fill-in events are mediated by Pol λ in human cells and that these insertions, but not those generated by other *POLL*-dependent processes, are suppressed by *XRCC5*, *XRCC6*, and *PRKDC*. To validate this observation, we measured the effect of *XRCC5* and *POLL* knockdown on insertion frequencies at five endogenous Cas9 target sites in a separate cell line, HeLa with dCas9-KRAB (Figures 4G and S5A). These experiments recapitulated our finding that knockdown of *XRCC5* specifically increases *POLL*-mediated duplication of PAM-distal nucleotides during repair of Cas9-induced DSBs.

*XRCC5* and *XRCC6* encode two subunits of a DSB-sensing heterodimer (Ku), which rapidly binds DSB ends and activates the transducing kinase DNA-PKcs (encoded by *PRKDC*) to coordinate the assembly and activation of many NHEJ factors^40, 44, 45^. Biochemical assays have demonstrated that Ku and DNA-PKcs can protect DNA ends from processing by DNA polymerases, including Pol λ^46^, and Ku facilitates error-free joining of cohesive ends^46, 47^. Synthesizing these observations with our screen results suggests that Pol λ-mediated processing is efficiently suppressed by Ku and DNA-PKcs at staggered cuts (Figure 4H) but not when Cas9 cuts with canonical blunt activity, indicative of a complex interplay between different repair factors and specific DSB configurations. More generally, these results highlight the ability of Repair-seq to identify diverse genetic dependencies behind outcomes with similar sequence architectures.

### Capture of human genomic sequence at DSBs is stimulated by loss of DNA2 and MCM10

In addition to short indels, we observed repair outcomes in which longer stretches of sequence from the human genome were “captured” at breaks^14, 48–50^ (6.6% +/- 2.5% baseline frequency across Cas9 screens, Figure S5B). While such outcomes were individually rare, analyzing changes in their collective frequency allowed identification of genetic determinants. Initial analysis revealed that the length distribution of captured genomic sequences in unperturbed cells consisted of partially-overlapping populations centered at ∼75 and ∼150 nts (Figure S5C). These two populations had dramatically different genetic dependencies, with a broad class of genes causing substantial changes in the shorter population but weak or no changes in the longer population (Figure S5D). This effect was most striking for two genes: (1) *DNA2*, which encodes a nuclease with high affinity for single-stranded 5’ flaps and multiple proposed roles in replication and DNA repair^51, 52^, and (2) *MCM10*, which encodes a replisome component with roles in replication initiation and elongation^53^ (Figure S5E-F). Knockdown of these genes produced 8- to 10-fold increases in the short population but substantially weaker changes in the long population. Notably, we also observed rare capture of sequence from the *Bos taurus* genome (0.1% +/- 0.06% of outcomes) (Figure S5G), presumably stemming from DNA in fetal bovine serum used in culture media entering cells during electroporation^54^. Similarly to capture of the longer population of human genomic sequences, capture of bovine sequences was not suppressed by knockdown of *DNA2* or *MCM10*, (Figures S5F and S5H), suggesting that the longer population of captured human sequences may also originate as extra-cellular DNA.

*DNA2* strongly suppressed capture of short segments of human genomic sequence at DSBs in our screens. Intriguingly, DNA2 deficiency was recently shown to increase the frequency of similar genomic capture events at induced breaks in yeast^50^, potentially by increasing levels of overreplicated single stranded DNA produced during Okazaki fragment maturation or the processing of reversed replication forks. Our results indicate that this phenotype is recapitulated in human cells. Furthermore, although our library targeted many different replisome components, screen results indicate that deficiency of one specific component, MCM10, also contributes as dramatically as DNA2 to this form of genomic instability. More broadly, identification of genetically distinct classes of genomic sequence capture events highlights how direct sequencing of outcomes allows Repair-seq to measure genetic determinants of a diverse range of repair events.

### Repair-seq allows systematic identification of complexes and pathways responsible for repair of Cas9- and Cas12a-induced DSBs

We next explored the idea that Repair-seq data could be used to group DSB genes into complexes and pathways. Hierarchical clustering of the outcome redistribution signatures produced by CRISPRi sgRNAs across all four Cas9 target sites revealed groups of sgRNAs that produced highly similar changes across outcomes, with several of these groups containing sets of sgRNAs targeting genes that encode subunits of known DSB repair complexes (Figure S4). A visually striking example grouped together sgRNAs targeting *XRCC5* and *XRCC6*. Another example included sgRNAs targeting MRN complex genes (*MRE11*, *NBN*, *RAD50*). Indeed, sgRNAs targeting genes associated with these complexes, as well as those encoding a third complex, replication factor C (*RFC2, RFC3, RFC4,* and *RFC5*), frequently correlated more highly with each other than with any other targeted genes (Figure 5A-5C).

**Figure 5.**
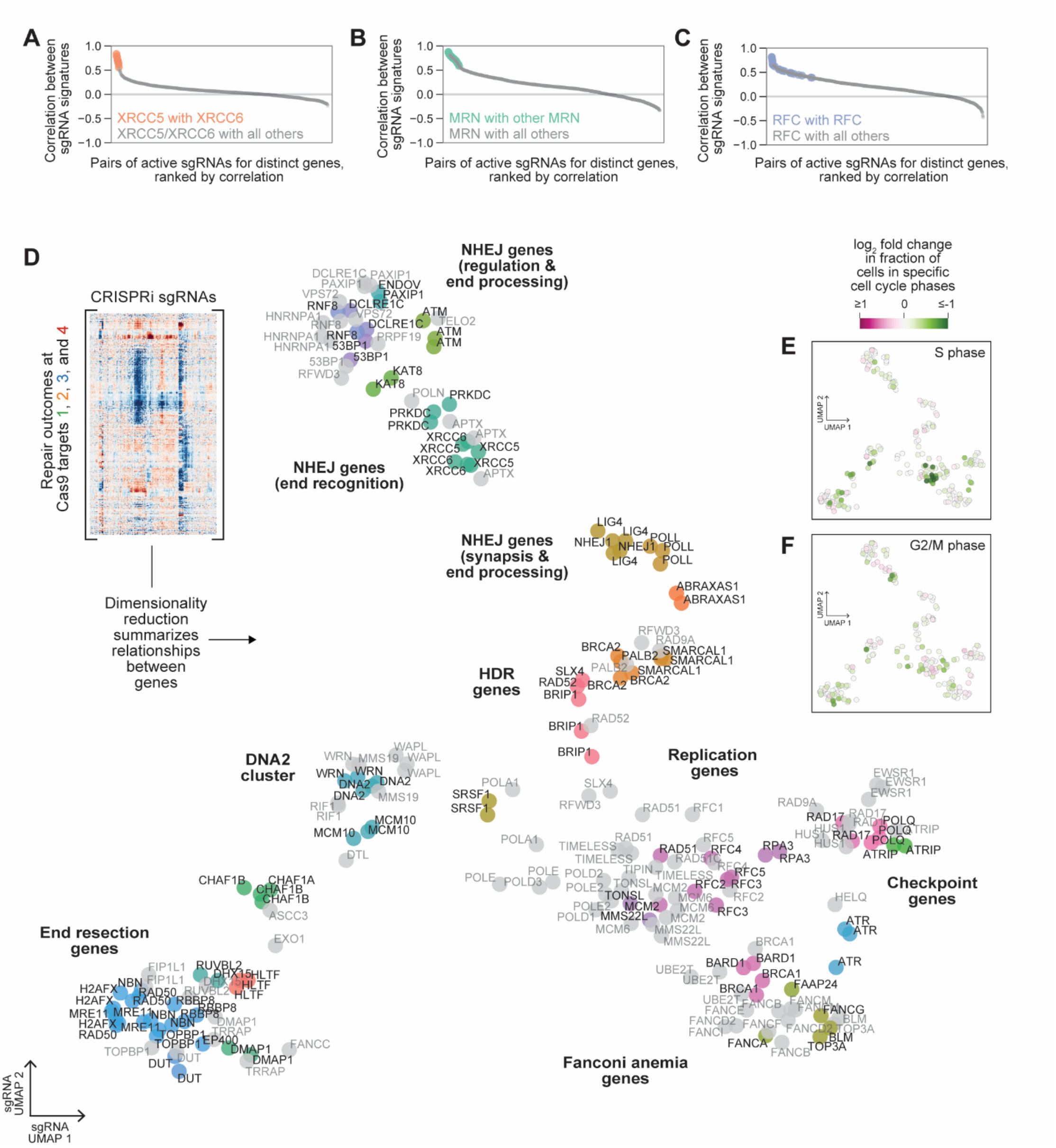
Systematic inference of functional relationships between DNA repair genes. (A+B+C) Correlations between outcome redistribution profiles for pairs of active sgRNAs targeting distinct genes where one or both genes is a member of a known complex (A: XRCC5/XRCC6; B: MRE11/RAD50/NBN, C: RFC2/RFC3/RFC4/RFC5). sgRNA pairs are plotted in rank order with colored dots marking pairs targeting distinct members of the complex. (D) sgRNA-centric dimensionality reduction process summarizes relationships between repair genes. Columns of the composite matrix of log_2_ fold changes in outcomes at four SpCas9 target sites for the most active CRISPRi sgRNAs (inset) are embedded into a two-dimensional space via UMAP to group different sgRNAs that modulate similar sets of outcomes. Points correspond to distinct sgRNAs and are colored by HDBSCAN assignment of clusters of sgRNA vectors prior to dimensionality reduction. (E+F) Changes in the fraction of cells that received each sgRNA assigned to S phase (E) or G2/M phase (F) relative to all non-targeting sgRNAs based on transcriptional profiles measured by Perturb-seq, overlaid on the sgRNA UMAP embedding of Repair-seq phenotypes.

To further explore gene-gene relationships with our data, we performed dimensionality reduction on the columns of our composite Cas9 outcome redistribution matrix (Figure S4), creating an embedding based on the outcome redistribution profiles of each sgRNA (Figure 5D). This embedding revealed many groups of genes with similar roles in DSB repair, including genes involved in DSB end resection (e.g., *MRE11*, *NBN*, *RAD50*, *RBBP8*), replication genes (e.g., *RFC1*, *RFC2*, *RFC3*, *RFC4*, *RFC5*, *MCM2*, *MCM6*, *TIMELESS*, *TONSL*, *MMS22L*, *POLE*, *POLE2*), replication checkpoint genes (e.g., *RAD17*, *HUS1*, *RAD9A*), HDR genes (e.g., *BRCA2*, *PALB2*), and Fanconi anemia pathway genes (e.g., *FANCI*, *FANCE*, *FANCD2*, *FANCB*, *FANCM*, *FANCF*, *FANCA*, *FAAP24*, *UBE2T*, *BRCA1*, and *BARD1*). This embedding also separated canonical NHEJ genes into multiple distinct clusters (e.g., *53BP1*, *RNF8*, *PAXIP1*, and *DCLRE1C; LIG4*, and *NHEJ1; XRCC5*, *XRCC6*, *PRKDC* and *APTX*), highlighting the power of high-resolution phenotypes to capture specific roles in DSB end joining. Moreover, consistent with the roles we identified for *MCM10* and *DNA2* in promoting genomic sequence capture events, these genes grouped together. Finally, this analysis nominated new roles for specific genes in DSB repair by forming unexpected groupings, including placing replication checkpoint genes together with *POLQ*, a regulator of alternative end joining, and placing *TOPBP1*, a canonical replication and checkpoint gene^55^, with those involved in end resection.

Because DSB repair is highly regulated by the cell cycle^56^, the effects of gene knockdown on cell cycle progression may contribute indirectly to observed changes in outcome distributions. Most notably, resection at DSB ends is largely restricted to cell cycle stages where a sister chromatid is present^56^. Perturbations that reduce occupancy of these stages would consequently be expected to decrease outcomes dependent on resection. We therefore measured the fraction of cells observed in S and G2/M phases for each CRISPRi sgRNA in our 366 sgRNA sub-library (after 6 days of gene depletion) using gene expression profiles collected via Perturb-seq, a method for performing single-cell RNA-sequencing of CRISPR-based screens^57^ (Method Details). Overlaying these phenotypes onto our sgRNA embedding revealed that sgRNAs with strong effects on relative occupancy of cell cycle phase were spread throughout the embedding, within and outside of distinct clusters (Figure 5E and 5F), indicating that outcome redistribution signatures of sgRNAs with cell cycle effects can provide additional signal. Clustering of *TOPBP1*, which has well-defined roles in replication initiation and checkpoint signaling, but which was grouped with end resection genes (i.e., *MRE11*, *NBN*, *RAD50*) (Figure 5D), provided an interesting example of this logic. The association between *TOPBP1* and genes involved in resection could be explained by the fact that repression of *TOPBP1* reduced the number of cells in S phase. However, we also found that other sgRNAs in the library, such as those targeting *MCM2*, caused stronger reductions in S phase occupancy but produced weaker correlations with the outcome changes produced by repressing *MRE11*, *NBN*, and *RAD50* (Figure S6A). Reduction in S phase occupancy produced by knockdown of *TOPBP1* thus may not be sufficient to explain the close phenocopying of MRN genes by *TOPBP1*. Intriguingly, TOPBP1 has been shown to physically interact with MRN^58, 59^.

We next asked whether the functional relationships identified in our Cas9 screens are specific to DSBs produced by Cas9 or represent more general features of DSB repair. Previous work^60–63^ and our results on the idiosyncratic processing of Cas9-induced DSB with short overhangs (discussed above) have demonstrated that DSBs formed with different end configurations can be repaired in different ways. To survey an additional end configuration, we next performed a Repair-seq screen using *Acidaminococcus sp.* Cas12a (H800A mutant) RNP and our 1,573 CRISPRi sgRNA library (Figure S1A).

DSBs produced by Cas12a differ from those produced by Cas9 in two key ways: Cas12a breaks consist of two staggered nicks that leave ∼5 nt 5’ overhangs and are located further from the PAM sequence^64, 65^. Consistent with these differences, data from our Cas12a screen revealed several Cas12a-specific distinct features. First, clustering of Cas12a outcomes based on their genetic signatures identified distinct groups of deletions spanning each of the two staggered nicks, with complex patterns of dependencies on different NHEJ factors between the groups (Figure S6B). Second, we observed a set of sgRNAs that specifically increased only a set of outcomes for which the PAM and protospacer remained unchanged (Figure S6B). Because these outcomes are susceptible to re-cutting^66^ but will only be observed when they do not acquire additional mutations, increases in these outcomes may be driven by sgRNA-induced reductions in the overall rate of cutting. Finally, baseline frequencies of capture of human genomic sequences at Cas12a- induced DSBs were 3.6-fold lower than in Cas9 screens (1.8% in Cas12a screen vs. 6.9% average across Cas9 screens), suggesting that such events favor Cas9 end configurations.

Despite these differences, clustering of sgRNA signatures produced with Cas12a recapitulated many of the gene-gene relationships observed with Cas9. We observed, for example, similar groups of NHEJ genes, including a group of end synapsis genes *LIG4* and *NHEJ1*, one containing *XRCC5* and *XRCC6* next to *PRKDC*, and one featuring *53BP1*, *RNF8*, *PAXIP1*, and *DCLRE1C* (Figure S6B). Additionally, as with Cas9, we observed a group containing *POLQ* together with replication checkpoint genes (e.g., *RAD17*, *HUS1*), and a separate group containing DSB end resection genes (e.g., *MRE11*, *NBN*, *RAD50*, *RBBP8*) that again included *TOPBP1*.

### Identification of genetically distinct alternative end joining mechanisms

A major goal in building Repair-seq was to systematically delineate DSB repair pathways. With this in mind, we were intrigued by the observation that *POLQ*, a polymerase responsible for a form of alternative end joining called microhomology-mediated end joining (MMEJ), clustered with genes that have no characterized roles in MMEJ (i.e., *RAD17*, *HUS1*, and *RAD9A*) and not with others that regulate a key step, DNA end resection (i.e., *MRE11*, *RAD50*, *NBN*, and *RBBP8*)^8, 67^ (Figures 5D and S6B). We reasoned that examination of the outcomes promoted by each set of genes could provide insight into alternative end joining.

First, to understand which set of outcomes drove the clustering of *POLQ* with *RAD17*, *HUS1*, and *RAD9A*, we compared the phenotypes of these genes. Examination across all Cas9-induced outcomes within our composite data set revealed that these genes closely phenocopied each other, albeit with different strengths (Figure S4). To quantify this similarity, we calculated the correlations between outcome redistribution profiles for all pairs of active sgRNAs targeting distinct genes within both our Cas9 and Cas12a datasets (Figure 6A). This analysis revealed that correlations between *POLQ* sgRNAs and multiple sgRNAs targeting *RAD9A, HUS1*, and *RAD17,* as well as one sgRNA targeting *RAD1,* were among the strongest observed for any pair of genes in our library. Using arrayed assays, we validated the similarity between *POLQ* and *RAD17* phenotypes at one endogenous target site in K562 cells (Figure S6C).

**Figure 6.**
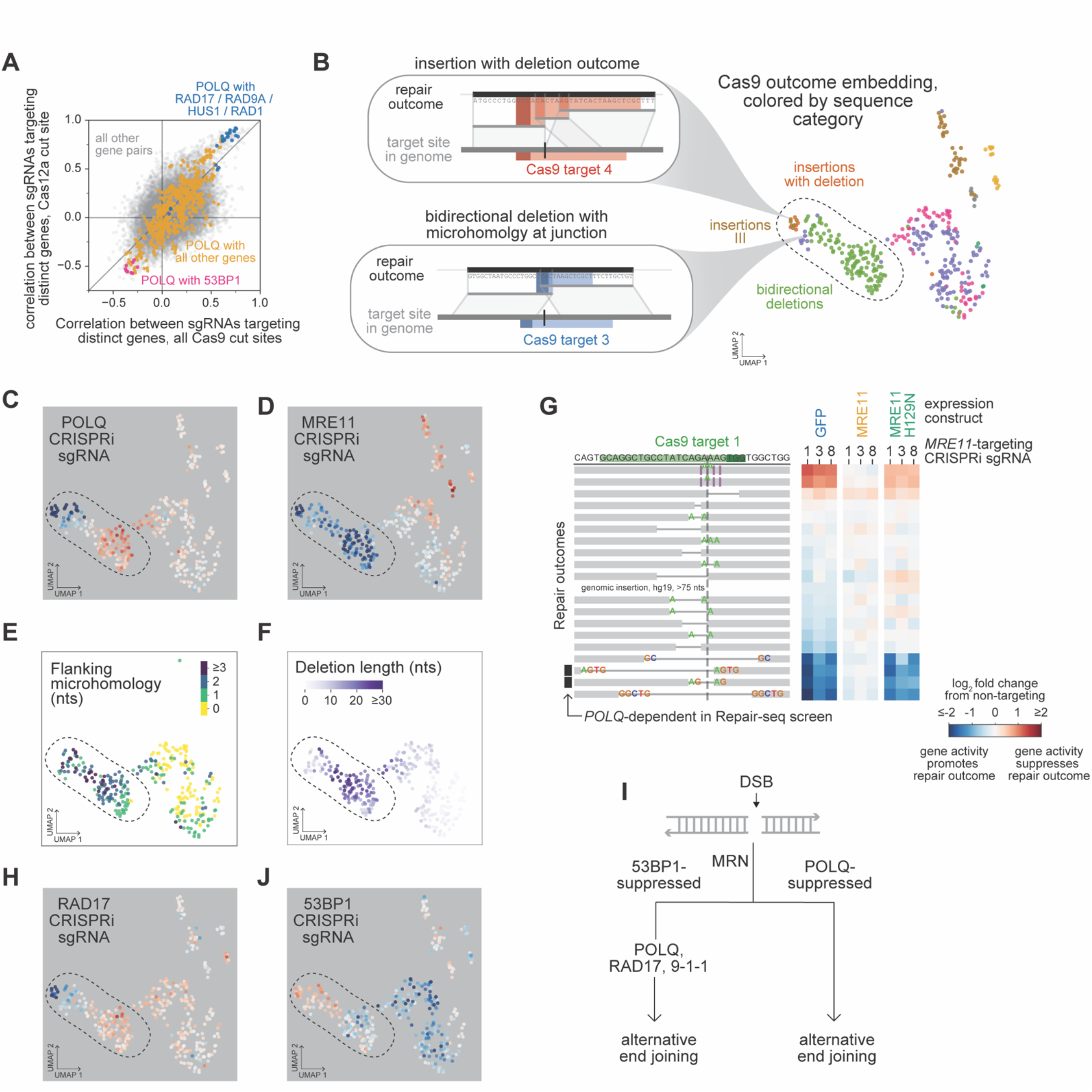
Repair-seq delineates alternative end joining mechanisms and nominates novel gene-gene functional relationships. (A) Comparison of correlations between outcome redistribution signatures for pairs of sgRNAs targeting distinct genes in composite SpCas9 screen data (x-axis) and Cas12a screen data (y-axis). Colors highlight specific indicated subsets of sgRNA pairs. (B) Outcome UMAP embedding for composite Cas9 data, with circled region containing outcomes consistent with known microhomology-mediated end joining (MMEJ) architectures: complex insertion+deletion sequence architectures (top inset), and bidirectional deletions flanked by microhomology (bottom inset). (C+D) Log_2_ fold changes in outcome frequencies produced by indicated CRISPRi sgRNAs overlaid on the composite Cas9 UMAP outcome embedding. (E+F) Deletions in the composite Cas9 outcome UMAP embedding, colored by the length of perfect microhomology flanking the deletion junction (E) or by the length of sequence removed by the deletion (F). (G) Effects of three *MRE11*-targeting sgRNAs on DSB repair outcomes in cells ectopically expressing GFP, MRE11, or nuclease-inactive MRE11 (H129N). Diagrams depict all outcomes present at baseline frequencies of at least 0.5%, sorted by the average phenotype across *MRE11* sgRNAs in cells with GFP. Heatmaps display log_2_ fold changes in outcome frequency for corresponding outcomes. (H) Log_2_ fold changes in outcome frequencies produced by a CRISPRi sgRNA targeting *RAD17* overlaid on the composite Cas9 UMAP outcome embedding. (I) Model of genetically distinct subpathways of alternative end joining. (J) Log_2_ fold changes in outcome frequencies produced by a CRISPRi sgRNA targeting *53BP1* overlaid on the composite Cas9 UMAP outcome embedding.

Next, we explored sequence characteristics of outcomes promoted by these genes. Previous work has defined two categories of mutations associated with Pol θ*-*mediated alternative end joining: (1) deletions of ∼5-50 nts with >2 nts of microhomology at the junction site^8^ (or within 15 nt on either side of the break site^4^, depending on the study), and (2) complex insertion outcomes in which DSBs are “filled-in” with sequences bearing microhomologies at the boundaries, potentially also accompanied by deletions^4, 68, 69^. The sequence characteristics of these outcomes are representative of Pol θ activity. Briefly, Pol θ is primed by short segments of complementary sequence, which arise at DSBs when base pairs on resected ends anneal across the break, or when other transient structures are formed^70^. DNA synthesis by Pol θ then either stabilizes the base pairing across the break to facilitate repair, leaving junctions that reflect characteristic microhomology usage, or adds templated base pairs to DSB ends, ultimately leading to insertions^71^. We observed examples of both such outcomes mediated by *POLQ* within our data set (Figure 6B and 6C).

End resection has also been identified as a key step in alternative end joining^8, 67^. Consistent with this, genes involved in end resection (i.e., *MRE11*, *RAD50, NBN*, and *RBBP8*) promoted a broad set of outcomes in our Cas9 screens that included those with expected architectures of Pol θ-mediated repair (outlined in Figures 6B, 6D, and S6D-F). This set of outcomes included insertions with microhomology at boundaries (Figure 6B and 6D; also discussed above as group III in Figure 4A), as well as a large set of longer, bidirectional deletions enriched for flanking microhomology (Figure 6B, 6D-6F). Examining *POLQ* dependency across these *MRE11*-dependent outcomes, however, separated them into two major groups: (1) a *POLQ-*dependent group containing insertions and a small subset of high-frequency *MRE11*- dependent deletions (Figures 3I, 6B, and 6C), and (2) a group of lower-frequency, typically longer *POLQ-* suppressed deletions (Figures 3I, 6B, 6C, and 6F). Together with similar observations from our Cas12a data (Figure S6B), these findings suggest delineation of two alternative end joining mechanisms downstream of *MRE11*-mediated end resection.

Given that MRE11, RAD50, NBN, and CtIP (encoded by *RBBP8*) have multiple roles in DSB repair, we next asked if *POLQ*-dependent and *POLQ*-suppressed deletions require the enzymatic activity of MRE11, which is responsible for nucleolytic processing of DSB ends. Consistent with a general requirement for resection across bidirectional deletions, treatment with the MRE11 exonuclease inhibitor mirin^72^ produced qualitatively similar outcome redistribution profiles as CRISPRi-mediated repression of *MRE11* at one endogenous target site in K562 cells (Figure S6C). Additionally, ectopic expression of nuclease active MRE11, but not a nuclease inactive mutant, alongside repression of endogenous *MRE11* rescued formation of deletions that both did and did not require POLQ in our screens (Figure 6G). These observations, together with the finding that phenotypes for *RAD17* and genes encoding components of the 9-1-1 complex (i.e., *RAD9A, HUS1*, *RAD1*) closely mirror *POLQ* phenotypes (Figures 6C, 6H, S4, and S6B), suggest that two alternative end joining mechanisms downstream of *MRE11*-mediated end processing contribute to DSB repair in K562 cells (Figure 6I). Moreover, the specificity of the relationship between *POLQ, RAD17*, and 9-1-1 genes, which promote and suppress the same subsets of *MRE11*- dependent outcomes in our screens, suggests a direct role for the products of those genes in promoting Pol θ-mediated repair, either through checkpoint signaling or via alternative functions.

Notably, a canonical characteristic of alternative end joining a lack of dependence on NHEJ factors^70^. The effects of NHEJ factors on *MRE11-* and *POLQ-*dependent outcomes within our maps, however, were not straightforward (Figures S4 and S6B). Loss of DNA-PKcs, for example, had varying effects across both *POLQ*-dependent and *POLQ-*suppressed outcomes in our Cas9 data set (Figure S6G). One exception, however, was *53BP1*, a factor that promotes NHEJ by suppressing resection^73, 74^. *53BP1* phenotypes clearly opposed those of *POLQ*, with specific suppression of the subset of *MRE11*-dependent, *POLQ*- promoted outcomes in both Cas9 and Cas12a datasets (Figures 6A, 6C, 6J and S6B).

### Repair-seq is a versatile platform for mapping the genetic determinants of genome editing modalities

Given our success mapping mechanisms of DSB repair, we next explored the flexibility of Repair-seq for broadly studying genome editing. Many genome editing approaches rely on endogenous DNA repair processes to install precise sequence changes^15, 75, 76^, and as demonstrated with our studies of Cas9 and Cas12a, Repair-seq can be easily tailored to different editing effectors. We therefore next applied Repair-seq to study single-strand template repair (SSTR) (Figure 7A), an exogenous form of HDR used for genome editing in which sequences from single-stranded oligodeoxyribonucleotides (ssODNs) are incorporated into nuclease-induced DSBs. Previous work has identified genetic regulators of SSTR in both yeast and human cells^7, 15, 77–80;^ however, the processes responsible for SSTR remain poorly defined.

**Figure 7.**
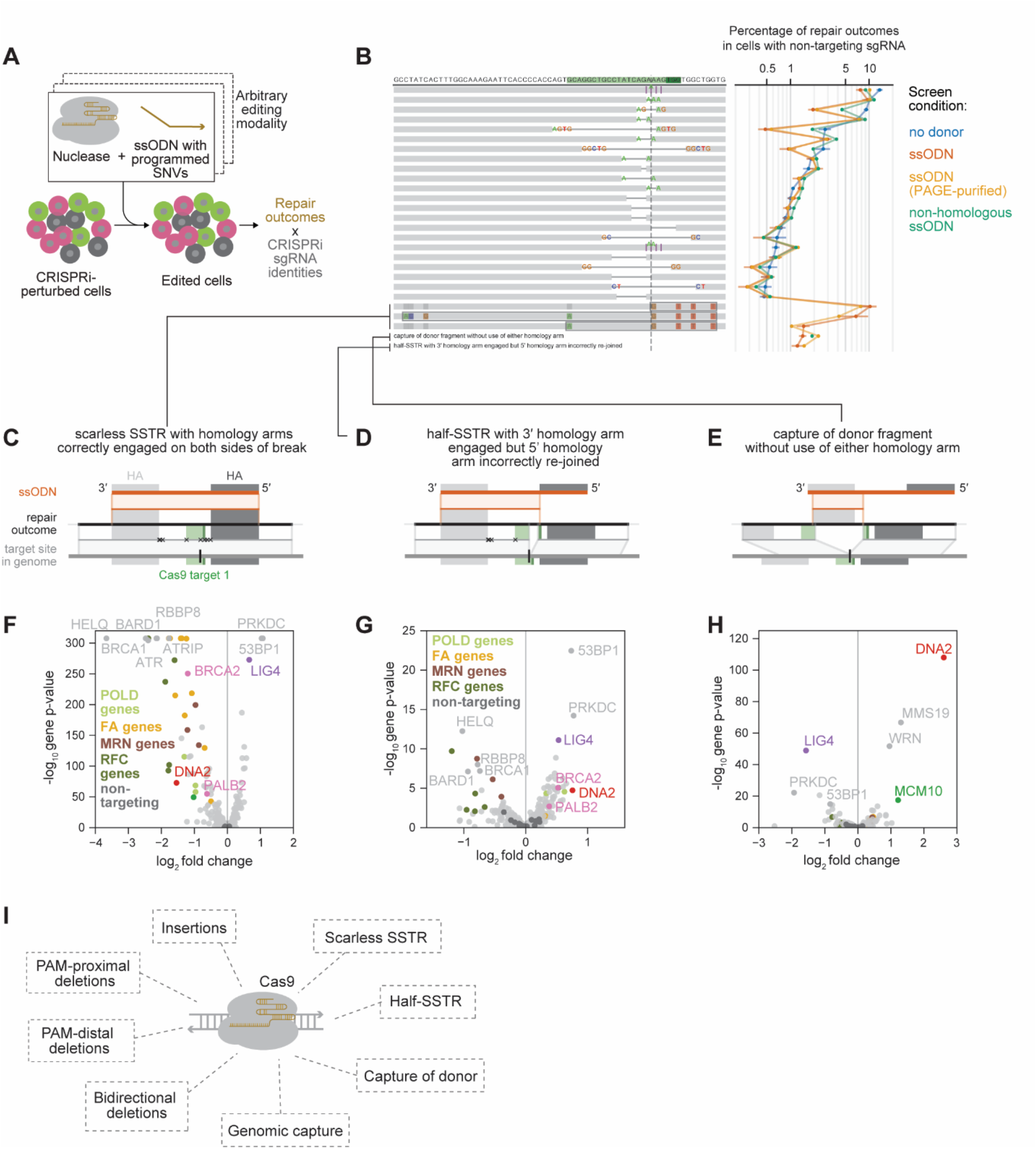
Repair-seq is a versatile platform for mapping the genetic determinants of diverse genome editing modalities. (A) Repair-seq can be adapted to study the effects of genetic perturbations on outcomes produced by diverse genome editing approaches. Electroporation of Cas9 RNP together with a single-stranded oligonucleotide DNA donor (ssODN) allows mapping of single-strand template repair (SSTR) mechanisms. (B) Comparison of baseline editing outcome frequencies observed in screens performed with and without donors: no donor (blue), an ssODN homologous to the target site (dark brown), the same ssODN stringently PAGE-purified (light brown), or an ssODN with no homology to the target site (green). To limit nuclease degradation, ssODNs had phosphorothioate bonds connecting the 3’ and 5’ terminal nucleotides. Diagram (left) depicts repair outcomes, sorted by average baseline frequency for non- targeting CRISPRi sgRNAs in no-donor screens. Plot (right) shows mean +/- standard deviation across replicates in baseline frequencies of each outcome in no-donor (3 replicates), homologous donor (4 replicates), and non-homologous donor (1 replicate) screens. Bottom two rows report the combined frequency of all outcomes with the sequence architectures shown in D and E. The three rows above these show scarless SSTR outcomes in which different subsets of donor-programmed SNVs have been installed. (C+D+E) Different categories of repair outcomes in which donor sequence has been incorporated. In each diagram, the center black line represents the sequencing read of a repair outcome, the top orange line represents the donor sequence, and the bottom grey line represents the sequence flanking the targeted DSB in the integrated screening vector. Lines between these three depict local alignments between the sequencing read and the relevant reference sequence, with any mismatch marked by an x. Grey rectangles mark homology arms (HAs) of matching sequence between the genome and the 5′ and 3′ ends of the donor. Green rectangles mark the protospacer and PAM of the Cas9 target site. Diagrams show examples of scarless HDR outcomes in which homology arms (HAs) are engaged on both sides of the break to drive scarless installation of any subset of donor-programmed SNVs (C), half HDR outcomes in which the 3′ HA is correctly engaged but the sequence outcome does not rejoin transition back to genomic sequence at the intended 5′ HA (D), and outcomes in which a fragment of donor sequence has been captured at the break without the use of either HA (E). (F+G+H) Volcano plots of the effects of gene knockdowns on the combined frequency of each donor- containing outcome category. Mean log_2_ fold changes in combined outcome frequency for the two most extreme sgRNAs targeting each gene (x-axis) and an aggregate binomial p-value for all sgRNAs targeting the gene (y-axis) are plotted. Black dots represent random sets of three individual non-targeting sgRNAs. Plots show effects on scarless SSTR outcomes (F), half-SSTR outcomes (G), and capture of donor fragments at the break (H). Gene categories indicated include DNA polymerase δ (POLD) genes, Fanconi anemia (FA) genes, MRN complex genes, and replication factor C (RFC) genes. (I) Cas9 induces diverse, genetically distinct mechanisms of repair of DSBs in the presence and absence of ssODNs.

To investigate SSTR, we performed Repair-seq screens using a set of ssODNs with homology to one of our Cas9 target sites, as well as a non-homologous control (Table S3). We started with a 178 nt ssODN containing 8 programmed single-nucleotide variants near the DSB, which were surrounded by homology arms (HAs) that perfectly matched the corresponding genomic sequence. Compared to screens performed without a donor, inclusion of the homologous donor changed baseline repair outcome frequencies in two informative ways. First, the frequencies of a specific set of indels enriched for bidirectional deletions with microhomology at the junction site were reduced (Figure 7B), suggesting that SSTR competes with specific donor-independent outcomes. The presence of a non-homologous donor produced more modest decreases in the same set of indels (Figure 7B). Second, we observed three major categories of outcomes in which donor sequence was incorporated at the break: SSTR outcomes in which different subsets of donor-encoded SNVs were scarlessly installed (Figure 7C); half-SSTR outcomes in which the HA on the 3’ end of the donor was correctly engaged, but the transition back to genomic sequence did not occur at the intended 5’ HA (Figure 7D); and capture of fragments of donor sequence into the break without using intended HAs on either side (Figure 7E). Arguing that contamination by truncated donor does not contribute substantially to formation of these outcomes, baseline frequencies of intended and unintended donor-containing outcomes were similar between screens performed with and without stringent purification of the donor (Figure 7B).

Comparing the genetic dependencies of each category of donor-containing repair outcomes revealed diverse mechanisms by which donor sequence can be incorporated at breaks. To identify regulators of scarless SSTR, we calculated the effect of gene knockdowns on the combined frequencies of all outcomes containing scarless installation of any subset of donor-encoded SNVs. Consistent with previous work, this bulk SSTR frequency was decreased by knockdown of many HDR genes (e.g., *BRCA1*, *RBBP8*, and Fanconi anemia genes) and increased by knockdown of NHEJ factors (e.g., *53BP1*, *PRKDC*, and *LIG4*) (Figure 7F). Intriguingly, knockdown of several genes that substantially reduced scarless SSTR did not reduce the frequency of half-SSTR outcomes, including *DNA2* and multiple genes involved in RAD51- mediated strand invasion (*BRCA2*, *PALB2*, *RAD51*, and *RAD51C*) (Figures 7F, 7G, and S7A). The differential effect of these genes on these two outcome categories indicates that their activities may play a specific role in pairing of the 5’ HA on the second side of the DSB. Finally, capture of partial donor fragments without the use of HAs was promoted by *LIG4* and strongly suppressed by *DNA2* but only modestly suppressed by *MCM10* (Figure 7H), suggesting that these events are mechanistically similar to capture of short human genomic sequences but with important differences (Figure S5E, see discussion). Capture of fragments of a non-homologous donor exhibited similar genetic dependencies (Figure S7B).

We next reasoned that examining the genetic dependencies of individual SSTR outcomes in which different subsets of SNVs were scarlessly installed could provide additional insights into SSTR mechanisms. To explore these outcomes, we performed screens using ssODNs containing 6 programmed SNVs evenly spaced around Cas9 target 1 with opposite strand orientations (i.e. whose sequences were the reverse complement of each other) (Figure S7C-S7E). Strikingly, across each of these donors, genes encoding components of the MutSα/MutLα DNA mismatch repair (MMR) complexes (*MSH2*, *MSH6*, *MLH1*, and *PMS2*) strongly promoted some scarless SSTR outcomes while suppressing others, with the suppressed set biased for outcomes missing SNVs from the 3’ ends of ssODNs (Figure S7C-S7E). These results are consistent with observations from previous studies, which proposed that MMR activity installs SNVs encoded on the 3’ ends of ssODNs by “fixing” mismatches established upon donor annealing^78, 81, 82^.

Our data extends this model by revealing a putative relationship between MMR genes and genes encoding components of the MRN complex (*MRE11*, *NBN1*, *RAD50*) (Figure S7C-S7E). Specifically, we observed two distinct levels of dependence on MRN components across SSTR outcomes, with outcomes exhibiting strong MRN dependence closely tracking outcomes suppressed by MMR, and outcomes exhibiting weaker MRN dependence closely tracking outcomes promoted by MMR. Together, these phenotypes suggest that incorporation of donor-encoded SNVs by MMR may relieve a requirement for end resection (Figure S7F). More generally, the existence of sets of SSTR outcomes with different levels of dependence on MRN highlight the diversity in mechanisms for arriving at donor-derived outcomes.

Collectively, these observations highlight the complex range of pathways by which ssODNs are incorporated at DSBs. Furthermore, our screens exploring SSTR mechanisms demonstrate that Repair-seq is well suited to delineating the genetic determinants of genome editing processes and identifying mechanisms that contribute to both intended and unintended editing outcomes.

## DISCUSSION

Here, we present Repair-seq, a combined experimental and analytical platform for systematic interrogation of DNA repair. By pairing CRISPR-based genetic screens with locus-specific deep-sequencing, Repair-seq allows massively parallel analysis of the effects of many genetic perturbations on the spectrum of DNA repair events following enzymatically induced changes to DNA. We applied Repair-seq to study repair of DSBs produced by 2 different enzymes (Cas9 and Cas12a) within 5 distinct sequence contexts in the absence and presence of homologous donors. Our efforts enabled a data-driven classification of DSB repair outcomes and their genetic dependencies. These classifications in turn produced a high-resolution atlas of repair mechanisms (Figure 7I), which recapitulates known repair pathways, initially defined through decades of focused studies, while also revealing multiple novel and mechanistically distinct repair processes.

A critical advantage of Repair-seq is the ability to classify DSB outcomes using genetic information without *a priori* registration of sequence features. Indeed, a recurring theme that arises from our studies is that repair outcomes with superficially similar sequence architectures can show marked differences in genetic dependencies. In one example, we identified an idiosyncratic role for the NHEJ factors Ku and DNA-PKcs in suppressing a set of Pol λ-mediated insertions. *In vitro* studies have shown that Ku and DNA-PKcs can promote non-mutagenic re-ligation of compatible ends by suppressing Pol λ fill-in^46^. Our observations indicate that this activity plays a major role in shaping repair outcomes at Cas9 breaks by specifically suppressing formation of insertions consistent with duplication of short Cas9-induced overhangs^42^, which are often the most frequent indel observed^12^. The counterintuitive fact that knockdown of core NHEJ factors leads to dramatic increases in this common NHEJ outcome highlights how complex interactions between repair factors and break configurations influence repair outcomes beyond simple pathway balance.

The ability to classify DSB repair outcomes with genetic signatures rather than assumptions based on sequence features also delineated pathways for alternative end joining that exploit short regions of homology to guide repair. We found that the canonical Pol θ-dependent MMEJ pathway in K562 cells is genetically distinct from an uncharacterized mechanism, which was suppressed by *POLQ* in our screens and requires *MRE11*-meditated end resection. This finding builds on a recent study that used targeted sequencing to classify alternative end joining outcomes^4^ and shows how, using Repair-seq, we can take otherwise uncharacterized sets of repair outcomes and assign them to discrete and novel pathways. Remarkably, from the same set of experiments, we identified previously unknown regulators of Pol θ- mediated MMEJ, including RAD17 and components of the associated 9-1-1 complex. We expect that our dataset, which will be available in a searchable online format, will act as a critical resource for future studies of alternative end joining.

Using Repair-seq, we also identified the genetic dependencies of more complex repair outcomes, uncovering a role for MCM10 in modulating capture of genomic sequence fragments at DSBs while also revealing important differences between its role and that of DNA2. We found that knockdown of either DNA or MCM10 dramatically increased the frequency of genomic sequence capture events in human cells. This DNA2 phenotype is consistent with recent observations in yeast^50^, in which DNA2 was proposed to suppress such outcomes by preventing generation of single-stranded DNA (ssDNA) through overreplication of genomic sequence fragments or by degrading intracellular ssDNA. In our screens, *DNA2* suppressed capture of exogenous ssODN fragments with similar magnitude as capture of genomic sequence fragments, but MCM10 knockdown had a much stronger impact on the latter. These results suggest that MCM10 deficiency has a specific role in generation of genomic fragments, while DNA2 activity downstream of fragment generation makes a substantial contribution to these phenotypes, either through degradation of single-stranded DNA or via more direct activity of the enzyme at DSBs. Together, these findings demonstrate how the ability to compare effects of genetic perturbations across multiple repair outcomes can both suggest and constrain models of the mechanisms behind repair events.

In the present study, we focused primarily on one cell type and growth regime, yet it is clear that many factors, including genetic background and local chromatin context impact DNA repair^27^. Applying Repair-seq to a diversity of dividing and nondividing cells, as well as exploring the effects of local transcription or chromatin modifications, will provide a powerful means to address open questions in DNA repair in a data driven and principled manner.

Finally, Repair-seq represents a flexible tool to understand and optimize diverse genome editing approaches. The rapid development of such approaches has dramatically outpaced our understanding of how different editing modalities engage with endogenous DNA repair mechanisms. We developed Repair-seq to investigate DSB repair, but the approach is readily adapted to study the repair of DNA lesions induced by alternative editing approaches. Critically, achieving precise editing requires understanding both the mechanisms that produce desired edits and the mechanisms that lead to unintended editing side products. The ability of Repair-seq to simultaneously interrogate the effect of genetic perturbations on a broad range of editing outcomes therefore makes it ideally suited to investigate editing tools. Our study of homology directed repair with single stranded oligonucleotide donors serves as a proof of principle for this concept, providing a systematic exploration of the diverse mechanisms by which donor sequence is incorporated at DSBs in both intended and unintended ways. Overall, the complexity of DNA repair mechanisms provides both a challenge and an opportunity for precision genome editing. The Repair-seq approach represents a powerful tool for unravelling this complexity.

## ACKNOWLEDGMENTS

We thank Marco Jost, Joseph Replogle, Manuel Leonetti, Hera Canaj, Bruce Conklin, and all members of the Adamson and Weissman labs for helpful discussions. We thank Michele Demozzi for identifying and screening gRNAs, Kenneth W. Gareau and Hayat S. Abdulkerim for RNP generation, and Chris Wilson and Mrudula Donepudi for reading the manuscript and providing helpful suggestions. We thank Eric Chow, Michelle Tan, Wei Wang, the UCSF Center for Advanced Technology, the Chan Zuckerberg Biohub, and the Genomics Core Facility of Princeton University for assistance with high-throughput sequencing. We thank Christina DeCoste and Katherine Rittenbach at the Princeton University Flow Cytometry Resource Facility, which is supported, in part, with funding from NCI-CCSG P30CA072720- 5921. Research reported in this publication was supported by the National Institutes of Health (NIH) under award numbers 1RM1HG009490 (J.S.W.), 1R35GM138167-01 (B.S.A.) and 5P30CA072720-22 (B.S.A.). J.S.W. is supported by HHMI. B.S.A. is supported by the Searle Scholars Program. J.A.H. was the Rebecca Ridley Kry Fellow of the Damon Runyon Cancer Research Foundation (DRG-2262-16). A.C. is supported by the National Science Foundation GRFP (DGE-2039656). A.C. and D.S. received support from a training grant from the NIH (T32HG003284). J.Y. is supported by a fellowship provided by the China Scholarship Council (CSC), based on the April 2015 Memorandum of Understanding between the CSC and Princeton University.

## AUTHOR CONTRIBUTIONS

J.A.H., C.C.R., J.S.W., and B.A. conceived of and designed Repair-seq. A.X. constructed the Repair-seq screening vector. A.X. and A.B. performed proof of principle experiments. A.X., A.C., and P.R. cloned the CRISPRi libraries. J.A.H., J.L., P.R., D.S., and B.A. performed Repair-seq screens, with assistance from A.X. and D.Y. J.Y. designed and performed ddPCR experiments. J.L., P.R., and B.A. performed validation experiments. J.A.H., A.C., and B.A. performed Perturb-seq. J.A.H. analyzed Repair-seq and Perturb-seq data. J.A.H., C.C.R., J.S.W., and B.A. interpreted screen results. J.A.H., C.C.R., J.S.W., and B.A. wrote the manuscript, with input from all authors. C.C.R., J.S.W., and B.A. supervised the project.

## RESOURCE AVAILABILITY

### Lead Contact

Further information and requests for resources and reagents should be directed to and will be fulfilled by the Lead Contact, Britt Adamson.

### Materials Availability

Plasmids and CRISPRi sgRNA libraries generated in this study will be made available on Addgene prior to publication.

### Data and Code Availability

Sequencing data will be made available on SRA prior to publication. Code is available at https://github.com/jeffhussmann/repair-seq, and data can be interactively explored at https://jeffhussmann.github.io/Repair-seq_demo/index.html.

## EXPERIMENTAL MODEL AND SUBJECT DETAILS

K562 cell lines were grown in RPMI 1640 with L-glutamine and 25 mM HEPES (Corning or Gibco) supplemented with 100 U/mL penicillin, 100 ug/mL streptomycin, and 0.292 mg/mL L-glutamine. 293T and HeLa cell lines were grown in DMEM with 4.5 g/L glucose and sodium pyruvate with or without L-glutamine (Corning or Gibco) supplemented with 100 U/mL penicillin and 100 ug/mL streptomycin. Cells were authenticated by analysis of short tandem repeats (ATCC) or by suppliers. One 293T cell line, used for packaging lentivirus, was replaced mid-study due to mutational events that reduced those cells to only a near perfect match to CCL-3216 (293-T). K562 cells expressing dCas9-KRAB from pHR-SFFV-dCas9- BFP-KRAB (Addgene, 46911) were previously described^24^ and were authenticated by analysis of short tandem repeats as an exact match to CCL-243 (K-562) from ATCC. HeLa cells expressing dCas9-KRAB from pHR-SFFV-dCas9-BFP-KRAB (Addgene, 46911) were previously described^29^ and were authenticated by analysis of short tandem repeats as an exact match to CCL-2 (HeLa) from ATCC. Mirin (Millipore Sigma, M9948-5MG) was dissolved in DMSO.

## METHOD DETAILS

### Plasmid construction

Our CRISPRi sgRNA libraries were delivered to cells using a custom expression vector (pAX198). We constructed this vector from pU6-sgRNA EF1Alpha-puro-T2A-BFP^24^ (Addgene, 60955) as follows: We added two restriction sites (NotI and HindIII) and spacer sequence to pU6-sgRNA EF1Alpha-puro-T2A-BFP to construct an intermediate vector (pMJ468). We then cloned a “target region” for enzyme-induced DNA damage between the sgRNA and selection marker expression cassettes (the latter expresses both BFP and puromycin resistance) by Gibson assembly at the NsiI and XhoI sites. This target region comprised sequence from the human *HBB* gene—specifically the second and third exons of ENST00000647020.1 (no intron) and part of the 3’ UTR—and was flanked by a NotI site. As part of this cloning, a second NotI site (adjacent to the target site) was removed and the termination signal for the U6- sgRNA expression cassette was reduced to a stretch of 6 thymines. Finally, to enable sgRNA cloning, a BstXI site was removed from the target site by site-directed mutagenesis. In arrayed experiments, individual sgRNAs were delivered using pU6-sgRNA EF1Alpha-puro-T2A-BF, including XRCC5_+_216974121.23-P1P2, POLL_-_103347937.23-P1P2, POLQ_+_121264772.23-P1P2, MRE11A_-_94227004.23-P1P2, RAD17_+_68665692.23-P1P2 (Table S2). pU6-sgRNA EF1Alpha-puro-T2A-BFP was used as a negative control in experiments depicted in Figures 4G and S5A and CRISPRi data in Figure S6C.

For the experiment depicted in Figure 6G, we built expression vectors using pInducer20 (Addgene, 44012) as follows: pInducer20 was first digested with AgeI and religated to remove unwanted functional cassettes. *EF1A* promoter was then inserted between the BamHI and XhoI sites. The resulting plasmid contains bicistronic expression cassettes with distinct promoters (*EF1A* and *UBC*) for each cassette. To generate *MRE11* expression vectors, superfolder GFP was inserted downstream of the *UBC* promoter between the KpnI and NotI sites, and *MRE11* cDNA sequence, amplified from pICE-HA-MRE11-WT (Addgene, 82033) using primer pairs oJL162 and oJL163 (Table S7), was inserted downstream of the *EF1A* promoter at the XhoI site. The nuclease-inactivating mutation (H129N) was then introduced by PCR with primer pairs oJL079 and oJL080 (Table S7). To generate a GFP expression vector control, superfolder GFP was inserted downstream of the *UBC* promoter between the KpnI and NotI sites. *MRE11*- targeting sgRNAs used in this experiment (1, 3, 8) were MRE11A_-_94227004.23-P1P2, MRE11A_-_94226994.23-P1P2, MRE11A_+_94226703.23-P1P2 (Tables S2 and S4). These sgRNAs and one non-targeting sgRNA (non-targeting_01332) were cloned into pAX198.

### Oligonucleotides

Oligonucleotides (unless otherwise indicated) and TaqMan probes were obtained from Integrated DNA Technologies (IDT). Single-strand oligonucleotides for SSTR experiments (oBA701, oJAH159, oJAH160, and oJAH165; Table S7) were ordered with phosphorothioate bonds connecting the 3’ and 5’ terminal nucleotides in order to limit nuclease degradation (represented by asterisks in Table S7). One single-stranded oligonucleotide (oBA701) was PAGE-purified by IDT or as follows: 25 µL of 100 µM oBA701 ultramer (10 nmol total) were combined with 25 uL 2x Novex TBE-Urea sample buffer (LC6876) and run on a 6% TBE-Urea gel (UreaGel System, National Diagnostics, EC-833). The gel was stained by washing 10X SYBR Gold (10000x concentrate, catalog number S11494, diluted in 1X TBE) over the surface of the gel and visualized using a Safe Imager 2.0 Blue Light Transilluminator (ThermoFisher Scientific). A high intensity band corresponding to the full length ultramer was cut out of the gel. Gel slices were pulverized using Gel Breaker tubes (Ist Engineering/Fisher NC0462125), and DNA was extracted from pulverized gel by soaking in 750 uL of extraction buffer (300 mM NaCl, 10 mM Tris 8, 1 mM EDTA) overnight. Extracted DNA was isopropanol precipitated, and the resulting pellet was cleaned up by ethanol precipitation.

### CRISPRi library design

For our 1,573 sgRNA CRISPRi library (AX227), we curated a set of 476 genes enriched for those involved in DNA repair and associated processes (e.g., DNA replication, repair, recombination) (Figure 1C and Table S5). We then selected 1,513 sgRNAs targeting these genes from a previously published CRISPRi library, hCRISPRi-v2.1 ^83^ (Table S2). Selected sgRNAs were those ranked as the top 3 per transcript per gene. A minority of sgRNAs were annotated as targeting multiple gene promoters. Annotations can be found in Table S2. We also included 60 non-targeting control sgRNAs. These were selected from hCRISPRi-v2^83^ randomly and filtered using data from a genome-scale growth screen in K562 cells^83^.

For our 366 sgRNA CRISPRi library (AC001), we selected 336 sgRNAs targeting 118 genes and 30 non-targeting controls (Table S4). To select the 118 genes targeted by this library (Figure S3B and Table S6), we considered data from Repair-seq screens performed with our 1,573 sgRNA CRISPRi library. Of the 118 genes, 114 are a subset of those targeted by our large library (AX227), while 4 were included based on interest from the literature (*EXD2*, *HELB*, *RBBP6*, *ZBTB38*). Each gene in this library was targeted with 2-6 sgRNAs (26 with 2, 86 with 3, 5 with 4, and 1 with 6). Among sgRNAs that target genes also included in AX227, we excluded 19 from AC001. A minority of sgRNAs included in AC001 have been annotated as targeting multiple gene promoters. Annotations can be found in Table S4.

### CRISPRi library cloning

Oligonucleotides containing sgRNA targeting sequences were synthesized by Twist Bioscience (Q-15620=AX227; Q-28859=AC001). These sequences were cloned into pAX198 with standard protocols (details available at https://weissmanlab.ucsf.edu/CRISPR/Pooled_CRISPR_Library_Cloning.pdf). Briefly, library sequences were amplified by PCR, purified (column-based), and digested with BlpI and BstXI. Digested fragments were then isolated by gel purification, precipitated using isopropanol (AC001) or ethanol (AX227), and ligated into a similarly digested vector (BlpI and BstXI). Ligations were performed at an insert to backbone ratio of 1:1 for 16 hours at 16 °C and subsequently electroporated into MegaX DH10B T1R Electrocomp™ cells (ThermoFisher Scientific). Cells were grown in liquid culture (∼1L) supplemented with carbenicillin (AC001) or on agar plates supplemented with carbenicillin and then scraped into liquid for plasmid purification (AX227). Final plasmid libraries were isolated using column-based purification. Activity of the 366 sgRNA CRISPRi library (AC001) was evaluated in K562 cells expressing dCas9-KRAB using Perturb-seq (Figure S3C and S3D).

### Perturb-seq screen

The relatively compact size of our 366 sgRNA CRISPRi library (AC001) allowed for characterization of transcriptional phenotypes produced by all sgRNAs in the library using direct capture Perturb-seq, a method for multiplexed single-cell RNA-sequencing of CRISPR-based screens^57^. To perform this experiment, we transduced the 366 sgRNA library (AC001) into K562 cells expressing dCas9-KRAB (9e6 cells) with centrifugation (2 hours at 1000 x g). Cells were grown for ∼6 days and selected with puromycin (3 µg/mL, starting on day 2). During selection, cells were split and media was replaced as needed. Selected cells were prepared for single-cell RNA-sequencing (scRNA-seq) as follows: Cells were centrifuged at 100 x g for 3 minutes to remove media, washed in PBS supplemented with 0.04% sterile-filtered Bovine Serum Albumin (Sigma Aldrich), centrifuged again at 100 x g for 3 minutes to remove wash, diluted with wash buffer, and then kept on ice before loading into droplets for scRNA-seq using the Chromium Single Cell 5’ Library & Gel Bead Kit (10x Genomics, PN-1000006) and the Chromium Controller (10x Genomics). Altogether, we performed 8 reactions of scRNA-seq (aiming for ∼14,000 cells per reaction). Cells were 88% viable and 94% BFP+ (sgRNA marker) before loading into droplets for scRNA-seq.

For scRNA-seq, we followed instructions in the Chromium Single Cell V(D)J Reagent Kits with Feature Barcode technology for Cell Surface Protein user guide (CG000186) with modifications to enable recording of sgRNAs as previously described^57^. Briefly, we added 5 pmol of a sgRNA-specific primer (oJR160, Table S7) to each reaction Master Mix prior to droplet formation. During cDNA amplification, we performed 11 cycles of PCR (according to manufacturer’s instructions) and after cDNA amplification separated the sample as follows: We performed a 0.6X left-sided cleanup reaction with SPRIselect (Beckman Coulter) and collected both the beads (which carry material for preparing the gene expression library) and the supernatant (containing sgRNA-derived cDNA amplicons). Using the supernatant, we then completed a 0.6X-1.35X double-sided cleanup. Eluate from the 0.6X cleanup was used to complete the gene expression library (according to manufacturer’s instructions), and eluate from the 0.6X-1.35X reaction was used to produce a library of sgRNA sequences containing scRNA-seq indexes.

Perturb-seq data from this experiment showed that sgRNAs in the 366 sgRNA CRISPRi library (AC001) produced a median of 86% depletion of target gene expression (Figure S3C), with 113 of 118 genes targeted by at least one sgRNA achieving >70% knockdown (Figure S3D). Additionally, using the Perturb-seq data, we quantified the effect of each sgRNA on relative occupancy of each cell cycle phase (Figure 5E and 5F).

### Ribonucleoprotein (RNP) complexes

*Streptococcus pyogenes* Cas9 and *Acidaminococcus sp. BV3L6* (A.s.) Cas12a with an RNase inactivating mutation (H800A) were complexed with guide RNAs obtained from Integrated DNA Technologies (Cas9, Alt-R® CRISPR-Cas9 crRNA and Alt-R® CRISPR-Cas9 tracrRNA; Cas12a, Alt-R® CRISPR-Cpf1 crRNA). Enzymes were obtained from IDT (Alt-R® spCas9 Nuclease V3, 1081059) or from Editas Medicine (made in-house or ordered from supervised vendor, Aldevron; sequences included below). RNPs obtained from Editas Medicine were complexed at a ratio of 2:1 (RNA:protein) as follows: Cas9 complexes were prepared to 50 µM by diluting Cas9 to 100 µM in complete 1X HG300 buffer (50 mM N-2-hydroxyethylpiperazine-N’-2-ethanesulfonic acid (HEPES), 300 mM NaCl, 1 mM tris(2-carboxyethyl)phosphine (TCEP), 20% glycerol (% v/v), pH 7.5) and mixing 1:1 with guide RNAs diluted to 200 µM in 1X H150 buffer (10 mM N-2-hydroxyethylpiperazine-N’-2-ethanesulfonic acid (HEPES), 150 mM NaCl, pH 7.5). HG300 buffer contained 300 mM of NaCl and H150 contained 150 mM of NaCl; the final salt concentration should therefore be 225 µM. Components were allowed to complex for at least 45 minutes at room temperature and a differential scanning fluorimetry assay^84^ was performed to confirm complexation occurred. Cas12 complexes were prepared to 66 µM by resuspending guide RNAs in H150:HG300 buffer to 200 µM and mixing with Cas12a (400 µM) and 1X H150 at a ratio of 4:1:1 (RNA:protein:buffer) by volume. Components were allowed to complex for at least 45 minutes at room temperature and a differential scanning fluorimetry assay was performed to confirm complexation occurred. Other RNPs (gRNA-5, gHEK3_3a, and gRNF2_2 in Table S1) were complexed as follows: crRNA and tracrRNA oligos were mixed in equimolar concentrations (1:1) to a final duplex concentration of 100 µM (16 µl total), incubated at 95 °C for 5 minutes, and allowed to cool to room temperature (15-25 °C) for ∼45 minutes. Duplexed guide RNAs were mixed with Cas9 enzyme and PBS (Thermo Fisher Scientific) to a final 12.2 µM RNP concentration and incubated at room temperature for ∼1 hr.

### Electroporation

For experiments depicted in Figures 4G, 6G, S1B-E, S2C, S2E, S5A and S6C, indicated cells were electroporated with RNPs as follows: Cells (1.25-6e5 or 3e6 per sample) were spun out of media by centrifugation (1000-1500 rpm for 5 minutes) and washed once with PBS (Thermo Fisher Scientific) or directly resuspended in electroporation buffer. Using an SE Cell Line 4D X Kit S (Lonza Bioscience), cells were electroporated on a 4D-Nucleofector (Lonza Bioscience) according to manufacturer’s instructions (CN-114 reagent code for HeLa cells and FF120 reagent code for K562 cells) using 50 pmol of Cas9 RNP. Details of electroporation during screens are described below (*Repair-seq Screens—cell culture*).

### Virus preparation

Lentivirus was produced in 293T cells by co-transfection of transfer plasmids (single or library) and packaging plasmids for expression of HIV-1 gag/pol and rev (+/-tat) and VSV-G envelope protein using either TransIT®-LT1 Transfection Reagent (Mirus) or Polyethylenimine (Polysciences, Inc.) with or without ViralBoost Reagent (Alstem, Inc.). For screening, virus-containing supernatant was either used directly, or filtered using either syringe filters (0.45 μm) or a vacuum filtration system (0.20 μm). Viral titers were determined by test transductions prior to screening (except for screen 1). For individual experiments, virus-containing supernatants were frozen prior to use.

### Repair-seq Screens—cell culture

K562 cells expressing dCas9-KRAB were transduced with an sgRNA library (AX227 or AC001) in large-scale infections (140e6 or 640e6 cells for AX227; 300e6 or 406e6 cells for AC001) (Table S3). Infections were supplemented with 8 μg/mL polybrene and conducted in many wells of multiple 6 well plates with centrifugation (∼2-3 hours at 1000 x g). After centrifugation, cells were pooled and resuspended in fresh, complete RPMI to ∼0.5e6 cells per mL. Screens 1-12 were spun out of virus-containing supernatant prior to resuspension. Cells were then grown with or without agitation for 6 days and selected with puromycin (added to 3 μg / mL beginning on day 2 for screens 1-5; added to 1 μg / mL beginning on day 2 and increased to 3 μg / mL on day 3 for screens 6-12; or added to 3 μg / mL beginning on day 3 for screens 13-18). For each screen, cells were less than 30% BFP+ (sgRNA library marker) two days post transduction. To remove dead cells, cultures were periodically spun out of media +/- PBS wash.

Screens were performed in the following groups (Table S3): 1; 2; 3-5; 6-12; 13-18. Screens that were performed concurrently were derived from a single transduction culture and split prior to RNP electroporation. Cells were electroporated with RNPs, with or without oligonucleotides, as follows: First, cells were spun out of media by centrifugation and washed with PBS. Next, using an SE Cell Line 4D X Kit L (Lonza Bioscience), cells were electroporated on a 4D-Nucleofector (Lonza Bioscience) with the FF120 reagent code, according to manufacturer’s instructions, with the following exceptions: (1) Post wash, cell pellets were mixed with Buffer SE (100 μL per reaction) and RNP (13 μL of 66 μM Cas12a or 5 μL of 50 μM Cas9 per reaction) with or without oligonucleotides (2.5 μL of 100 μM ssODN). (2) These mixtures were then loaded into cuvettes, typically using as many cuvettes as necessary to process the entire volume of cell mixture (∼100 μl per cuvette; numbers of cuvettes per screen indicated in Table S3). All RNPs used in screening were prepared and provided by Editas Medicine. Following nucleofection, cells were allowed to recover for ∼10 minutes at 37 °C (except for screen 11). Cells were then resuspended with media and transferred for growth. For screens 1 and 13-18, cells were spun out of media and washed with PBS two days post electroporation to help remove dead cells. After 3 days, cells were collected and cell pellets were processed to generate sequencing libraries (numbers of cells collected per screen, viability of cells at collection, and cell population doublings between nucleofection and collection are indicated in Table S3). Throughout screening, cells were counted and viabilities were determined using an Accuri™ C6 flow cytometer (BD Biosciences) or Countess II Automated Cell Counter (Thermofisher Scientific). Cells were typically maintained at densities between ∼0.5e6 and ∼1e6 cells per mL (splitting as necessary).

### Repair-seq Screens—sequencing library preparation

Sequencing libraries were prepared from the cells collected at the end of the screens as follows: Genomic DNA was extracted using the NucleoSpin® Blood XL kit (Macherey-Nagel) with a condition of 50e6-100e6 cells per column. Genomic DNA was then digested with NotI-HF (NEB) and run on large 0.8% agarose gels (Owl^TM^ A1 Large Gel System, Thermo Fisher Scientific) prepared with custom 3D printed combs capable of producing wells large enough for loading 1.5 mL volume per well (for more information on these gels, see https://weissmanlab.ucsf.edu/CRISPR/IlluminaSequencingSamplePrep_old.pdf), with individual samples typically spread across multiple wells. Fragments of genomic DNA within the size range of those containing both edited sequences and sgRNA expression cassettes (target fragments) were then excised with wide boundaries, purified using NucleoSpin® Gel and PCR Clean-up kit (Macherey-Nagel), digested with HindIII-HF (NEB) to remove a NotI overhang from one end of the target fragments, and isolated using SPRIselect Reagent (Beckman Coulter) in a 0.8X reaction.

Custom adapters containing 12 nt unique molecular identifiers (UMIs) were then ligated onto target fragments using the remaining 4 nt NotI overhang. These adapters (oBA676 and oBA677; Table S7) were obtained as individual DNA oligonucleotides (HPLC purified) from Integrated DNA Technologies and then annealed. For the ligation reactions, we used enzyme and buffer from the KAPA HyperPrep Kit (Roche). Ligation reactions were assembled as follows: 30 µL ligation buffer, 10 µL ligase, adapter at 200:1 adaptor:insert ratio, 1 µg of HindIII digested product, and PCR-grade water to 110 µL total volume. These reactions were incubated at 4 °C overnight on a thermocycler with lid temperature set to 30 °C. Following ligation, DNA was purified using SPRIselect Reagent (Beckman Coulter) in two steps: First, we performed a 0.65X reaction as calculated; however, as the ligation buffer contains PEG, the reaction ratio was effectively higher. Then, we performed a 0.8X reaction.

To enrich our target fragments for sequencing, we next amplified those sequences by PCR, performing enough PCR reactions to use nearly the entirety of each sample obtained from the ligation and subsequent clean-up reactions (number of PCR reactions per screen indicated in Table S3). We assembled PCR reactions with 30-50 ng of template into 50 µL total volume. These reactions contained amplification primers at 0.6 µM final concentration (each), 3% dimethyl sulfoxide, and 1X KAPA HiFi HotStart ReadyMix and were run on a thermocycler with the following program: 1 cycle of 3 minutes at 95 °C; 16 or 20 cycles of 15 seconds at 98 °C, followed by 15 seconds at 70 °C; 1 cycle of 1 minute at 72 °C; 4 °C hold. Amplification primers used were Illumina P7, which is included in the adapter, and an indexed version of ScreenIndexing (Table S7). Amplified DNA was purified for sequencing using SPRIselect Reagent (Beckman Coulter) in a 0.8X reaction, and index samples were mixed. Throughout the sample preparation, samples were checked for quality and yield using either a NanoDrop Spectrophotometers (Thermo Fisher Scientific), Agilent 2100 Bioanalyzer system, or by running on a Novex™ TBE Gel.

### Arrayed DSB repair experiments

For arrayed experiments depicted in Figures S1B-S1E, 4G, and S5A, as well as CRISPRi experiment in Figure S6C, cells were electroporated with indicated RNPs (as described above). Three days post electroporation, cells were collected and cell pellets were processed to generate sequencing libraries as follows. Genomic DNA was extracted using the NucleoSpin® Blood Mini kit (Macherey-Nagel). The targeted loci were amplified with primers (oJAH262 and oJAH263 for HEK3_3a target site, oJAH264 and oJAH265 for RNF2 target site, oJAH258 and oJAH259 for Cas9 target sites 1, 3, and 4 at the endogenous locus, oBA991 and oBA992 for Cas9 target sites 1, 3, and 4 within a genomically integrated screening construct; Tables S1 and S7) using NEBNext Ultra II Q5 Master Mix (New England Biolabs). Each reaction (100 µL total volume) contained 1 µg of genomic DNA and 1 µM of each primer and was run on a thermocycler with the following program: 1 cycle of 30 seconds at 98 °C; 22 cycles of 10 seconds at 98 °C, followed by 75 seconds at 65 °C; 1 cycle of 5 minute at 65 °C; 4 °C hold. The resulting products were purified using SPRIselect Reagent (Beckman Coulter) in 1X to 1.8X reactions and used as templates for a second PCR to add indexes to the amplified fragments. The indexing primers used were versions of Indexing_REV and Indexing_FOR in Table S7. Each of these reactions (50 µL total volume) contained 10 ng of template DNA and 0.6 µM of each primer with 1x KAPA HiFi HotStart Ready Mix and was run on a thermocycler with the following program: 1 cycle of 3 minutes at 95 °C or 98 °C; 8 cycles of 20 seconds at 98 °C, followed by 15 seconds at 65 °C and 15 seconds at 72 °C; 1 cycle of 1 minute at 72 °C; 4 °C hold. Amplified DNA was purified for sequencing using SPRIselect Reagent (Beckman Coulter) in 1X to 1.8X reactions, and index samples were mixed prior to sequencing. For experiment depicted in Figure S1F-S1E, K562 cells transduced with pAX198 were electroporated with RNPs against target sites 1, 3, and 4 (Table S1) (as described above). For experiments depicted in Figures 4G and S5A, as well as CRISPRi experiment in Figure S6C, K562 cells expressing dCas9-KRAB or HeLa cells expressing dCas9-KRAB were transduced with indicated sgRNA and selected with puromycin (1-3 µg/mL) prior to electroporation.

For mirin experiment depicted in Figure S6C, K562 cells were electroporated with Cas9 RNP against target site 1 (Table S1) and treated with mirin (Millipore Sigma, M9948-5MG) 20 minutes after electroporation at 50 µM final concentration. After 2 days, cells were collected and processed to generate sequencing libraries. Each sample was lysed with 500 μL lysis buffer (10 mM Tris-HCl, pH 7.5, 0.05% SDS, and 25 μg/mL Proteinase K) at 37 °C, followed by 30 minutes at 80 °C, and the lysate was used for PCR directly. The targeted locus was amplified using oJAH258 and oJAH259 with Phusion U Green Multiplex PCR Master Mix. Each reaction (25 µL total volume) contained 1 μL of template and 0.5 µM of each primer and was run on a thermocycler with the following program: 1 cycle of 2 minutes at 98 °C; 30 cycles of 10 seconds at 98 °C, followed by 20 seconds at 65 °C and 30 seconds at 72 °C; 1 cycle of 2 minute at 72 °C; 4 °C hold. The amplified fragments were indexed with a second PCR as described directly above, except that 10 ng templates were used in the reaction and 10 PCR cycles were run instead of 8. Indexed samples were purified using SPRIselect Reagent in a 1X reaction and pooled for sequencing.

To control for potential differences in overall rates of DSB induction between different arrayed experiments, repair outcomes reporting the unmodified sequence were excluded from analysis of arrayed experiments.

### MRE11 rescue experiment

For the experiment depicted in Figure 6G, constructs expressing GFP, *MRE11*, and a nuclease-inactive mutant (MRE11-H129N) were transduced into K562 cells expressing dCas9-KRAB alongside a single CRISPRi guide (one of three *MRE11*-targeting sgRNAs or non- targeting_01332). Cells infected with the same expression construct (different sgRNAs) were pooled together and, after 3 days, cells expressing GFP (expression construct marker) and BFP (sgRNA marker) were isolated by fluorescence-activated cell sorting on a FACSAria Fusion (BD Biosciences). On day 6, cells were electroporated with RNP (Cas9 against target site 1) as described above. After 3 days, cells were collected and cell pellets were processed to generate sequencing libraries as described above for arrayed experiments, except that 1.8 µg genomic DNA was used for the first PCR amplification. Notably, all of the *MRE11*-targeting sgRNAs used in this experiment were designed to target endogenous *MRE11* and are not expected to deplete exogenous MRE11 expressed from the *EF1A* promoter.

### Sequencing

Sequencing libraries from Repair-seq screens were sequenced on an Illumina NovaSeq 6000 System with 4 total reads per cluster (paired end reads plus two additional indexing reads) with the following read lengths: I1 = 12 nts (UMI sequence); I2 = 8 nts (sample index); R1 = 45 nts (CRISPRi sgRNA); R2 = 258 nts (editing outcome) (Figure S1F). Reads were demultiplexed based on sample index and CRISPRi sgRNA, allowing up to one mismatch between observed sequences and expected sample indices or spacer sequences. Reads reporting a CRISRPi spacer sequence of GACCAGGATGGGCACCACCC represented delivery of the sgRNA present in pAX198 before sgRNA library cloning; these reads were disregarded. Demultiplexing code can be found in https://github.com/jeffhussmann/repair-seq/blob/master/demux.py. Across screens, we obtained ∼0.03 UMIs per cell collected at the end of the screen. Number of sequencing reads, total UMIs, and median UMIs per guide obtained from each screen are indicated in Table S3. Sequencing for Perturb-seq experiment was performed on an Illumina NovaSeq 6000 system with paired end reads (I1 = 8nts; R1 = 28 nts; R2 = 98 nts). Sequencing of arrayed experiments and *MRE11* rescue were performed on an Illumina MiSeq System with single end reads (I1 = 8 nts; I2 = 8 nts; R1 = 300 nts) or paired end reads (I1 = 8 nts; I2 = 8 nts; R1 = 44 nts; R2 = 240 nts).

### Digital Droplet PCR (ddPCR) to detect large deletions

Some DSB repair products can remain “unseen” when analyzed by short-read sequencing. We observed that, after RNP electroporation, a fraction of electroporated cells lost expression of a BFP marker encoded on our screening construct (∼2000 nt away from the target region) (Figure S2A). Further investigation of this phenomenon by ddPCR analysis showed that BFP- populations isolated by fluorescence-activated cell sorting (FACS) were enriched for cells lacking sequence within a 4069 nt window around the targeted DSB, as compared to BFP+ populations isolated from the same electroporation reaction (Figure S2B and S2C). This observation is consistent with reports of large deletions induced by endonuclease cutting ^33^; however, we also observed that some of these repair products represented repair junctions wherein the flanking lentiviral long terminal repeats (LTRs) collapsed, indicative of repair through single-strand annealing (SSA) (Figure S2D and S2E). Given the relative positions of sequences important for Repair-seq sequencing library preparation (Figure S2B), these events would not be detected.

For the experiments in Figure S2B-S2E (described immediately above), we constructed 3 cell lines. First, we transduced K562 cells expressing dCas9-KRAB with pAX198 and selected them with puromycin. Cells expressing BFP were then isolated by FACS (FACSAria Fusion, BD Biosciences) to establish (1) a BFP+ cell pool and (2) two clonal BFP+ lines. To ensure homogeneous expression of BFP, both clonal lines were checked by flow cytometry (LSR II Flow Cytometer, BD Biosciences), and we verified single-copy lentivirus integration for each by ddPCR (QX200 AutoDG Droplet Digital PCR system, Bio-Rad). For clonal line 2, we determined the lentiviral integration site using a protocol adapted from Lenti-X Integration Site Analysis Kit (Takara Bio) and oligonucleotides oJY0104-oJY0109.

ddPCR experiments were then performed as follows: First, our established BFP+ cell pool and/or BFP+ clonal line(s) were electroporated with Cas9 RNP against target site 1 (Table S1) or Cas9 programmed with a negative control gRNA (gRNA-neg in Table S1) as described above. Four days post nucleofection, BFP+ and BFP- cell populations were separated by FACS (FACSAria Fusion, BD Biosciences). Genomic DNA was then extracted from each population using the NucleoSpin Blood, Mini kit for DNA from blood (Macherey-Nagel) according to manufacturer’s instructions. ddPCR was performed using this DNA, a forward primer, a reverse primer, and a TaqMan probe for each amplicon indicated in Figure S2B and S2D, as well as an *ACTB* control amplicon (oBA074, oBA075, oJY012, oJY0117, oJY033, oJY0034, oJY0037-39, oJY0089, oJY0090, and oJY040; Table S7). Amplicons used to quantify large deletions surrounding the edit site were 298 nt upstream, 126 nt downstream, and 3453 nt downstream of the DSB (designed using Primer3Plus; Figure S2B). For our SSA assay, we designed two amplicons specific to clonal line 2, using the sequence of the lentiviral integration site: (1) the “LTR amplicon” (designed using Primer3Plus), which was amplified by primers that bound immediately upstream and downstream of the 5′ LTR (oJY0110, oJY0115; Table S7), and (2) the “SSA amplicon”, which was amplified by oJY0110 and a primer that binds immediately downstream of the 3′ LTR (oJY0113; Table S7). A single TaqMan probe (oJY0099; Table S7) was used for both amplicons. For each amplicon, raw concentration measurements were normalized to the *ACTB* control amplicon from the same sample. Comparison of the normalized concentration of the “SSA amplicon” to that of the “LTR amplicon” in cells electroporated with Cas9 and gRNA-neg (Table S1) allowed quantification of the former. ddPCR reactions were performed using Bio-Rad ddPCR Supermix for Probes (no dUTP) following manufacturer’s instructions with a QX200 AutoDG Droplet Digital PCR system.

### Flow cytometry

Flow cytometry data was collected with an LSR II Flow Cytometer (BD Biosicences) and FACSDiva™ Software (BD Biosicences). Data in Figure S2A was processed with FCS Express 7 Research software (Version 7.04.0014).

## QUANTIFICATION AND STATISTICAL ANALYSIS

### UMI consensus calling

Within each sample index and CRISPRi sgRNA, reads are grouped by UMI sequence. Distinct reads with the same sample index, CRISPRi sgRNA, and UMI may represent amplification products from the same initial UMI-ligated molecule (possibly modified by point mutations or short indels from PCR or sequencing errors) but may also represent chimeric products generated by recombination during PCR or by index switching during cluster generation on Illumina platforms that use exclusion amplification for cluster generation^85^. The goals of processing each group of reads with the same UMI are to cluster together reads likely to represent the same repair event and use the redundant information present across such reads to correct sequencing or PCR errors by identifying a consensus sequence, and to filter out reads likely to represent recombination/index switching artifacts.

To accomplish these goals, a greedy clustering algorithm consisting of the following iterative process is applied to each UMI read group. First, the most common sequence in the group is identified. Then, all reads for which no base calls with quality score greater than 20 disagreed with the identified common sequence are removed from the read group to form a cluster. A consensus sequence of the cluster is called by identifying the most frequent base call across all reads at each position. If more than one base call is equally frequent, the consensus base is set to ‘N’ and consensus quality score is set to 2. Positions at which a single base call is present in more than 50% of reads for which at least one such base call had a quality score greater than or equal to 30 are assigned a consensus quality score of 31. All other positions are assigned a consensus quality score of 10. This consensus formation process is applied to both the repair outcome sequence and the CRISPRi sgRNA sequence. This process is iteratively repeated on remaining reads in the UMI read group until all reads have been assigned to clusters. Any cluster containing fewer than 4 reads is discarded. Of the remaining clusters, the cluster containing the highest number of reads is retained and all others are discarded. Finally, clusters for which the consensus CRISPRi sgRNA sequence did not perfectly match the designed protospacer may represent sgRNAs that experienced an error during synthesis, cloning, or lentiviral packaging. Since such errors may result in weaker knockdown of the targeted gene, any such clusters are discarded. All remaining clusters are advanced to outcome categorization.

### Categorization of repair outcome sequences

Consensus sequences of editing outcomes are classified using a modified version of *knock-knock*. Conceptually, the process of classifying a given sequence outcome consists of first generating a comprehensive set of local alignments between the outcome sequence and all relevant references sequences: the lentiviral screen vector, any donor sequences supplied in the screen, the human (hg19) genome, and the cow (bosTau7) genome. Alignments to the screen vector and donor sequences are generated using BLASTN version 2.10.1+ (with parameters described in https://github.com/jeffhussmann/knock-knock/blob/master/knock_knock/blast.py) and using a custom Smith-Waterman implementation (https://github.com/jeffhussmann/hits/blob/master/hits/sw.py). Alignments to genomes are generated using STAR version 2.6.0a. All local alignments for a given read are then jointly processed to identify the arrangement of segments of the screen vector and any other relevant sequence sources that make up the outcome and to identify any mismatches, insertion, deletions, or other rearrangements of these sequences to assign a classification to the type of editing outcome represented by the read. A full delineation of the decision tree carried out during this classification process can be found in https://github.com/jeffhussmann/repair-seq/blob/master/pooled_layout.py.

A summary of the portions of the decision tree relevant to results presented in the manuscript is presented below. The categorization process assigns a “category”, “subcategory”, and “details” to each outcome. The process first attempts to identify a single alignment to the screen vector, potentially containing a single deletion or short insertion up to 2 nts but not containing any mismatches, that begins by pairing the beginning of the read with the expected sequencing primer and that extends to the end of the sequencing read. If such an alignment exists with no insertion or deletion, the outcome is classified as category “wild type”. If the alignment contains an insertion, the outcome is classified as category “insertion” and subcategory “insertion”, with the inserted sequence and the position in the target after which the insertion begins recorded as details. If the alignment contains a deletion, the outcome is classified as category “deletion” and subcategory “near cut” if the deleted region overlaps a window of 5 nts around the expected cut site and subcategory “far from cut” if it does not, with the length of the deletion and the position in the target at which it begins recorded as details. The exact insertion or deletion assigned by the alignment process may represent an arbitrary representative from a class of equivalent insertions or deletions that would have produced indistinguishable resulting sequences. To account for this, the screen vector sequence is preprocessed to identify classes of degenerate insertions (that is, distinct combinations of inserted sequence plus insertion location that result in identical post-insertion sequences) and degenerate deletions (distinct starting locations of a given deletion length that result in identical post-deletion sequences). A lookup table from each member of a degenerate class to the representation of the full class is populated, and the observed insertion or deletion is looked up in this table to possibly replace its details with the details of the degenerate class of which it is a member.

If a single read-spanning alignment contains mismatches relative to the screen vector, these mismatches are compared to the expected SNVs programmed by a donor sequence. If all mismatches correspond to donor SNVs, the alignment contains no insertions, and the alignment contains no deletions longer than 1 nt long, the outcome is assigned category “HDR”. If there are no deletions, the outcome is assigned subcategory “clean”. Empirically, in reads that contain donor SNVs, we frequently observe 1 nt deletions throughout the region corresponding to the donor sequence. Deletion of 1 nt is a common error mode of oligonucleotide synthesis, so we conservatively attribute such deletions as likely existing in the starting oligonucleotide donors rather than being introduced by an error-prone repair process. If there are 1 or more 1 nt long deletions in the alignment, the outcome is therefore assigned subcategory “synthesis errors”. These outcomes correspond to the scarless SSTR category in Figures 7 and S7.

If no single alignment to the target covering the whole read is identified, the longest alignments to the target that reach each edge of the read are identified. Note that while the start of the read should always be covered by such an alignment because of the amplification strategy, the end of the read may not be covered if a sufficiently long sequence fragment has been captured at the DSB. The gap between these target edge alignments on the read is determined, and the remaining local alignments are searched to attempt to explain the gap. If no gap-covering alignments exist, a more permissive application of Smith-Waterman is applied to specifically search for additional local alignments between the gap region and the screen vector or the sequence of any donor used.

If the gap can be covered by an alignment to donor sequence, the outcome is assigned category “donor misintegration”. Donor misintegrations are sub-categorized based on how the donor sequence lines up with the target edge alignments on each side of the cut site. The two sides of the cut site relative to the read (left and right) are separately classified. For each side, if the homology arm in the donor alignment and in the target edge alignment are aligned to the same portion of the read, that side is classified as “intended”. If this is not true and if the donor alignment on a side extends all the way to an edge of the donor, the side is classified as “blunt”. If neither of these conditions is true, the side is classified as “unintended”. An additional wrinkle exists on the right side of the read, where the sequencing read may not be long enough to reach the end of the donor alignment and therefore may not be able to resolve how the outcome transitioned from donor sequence back to target sequence on the right side. If this is the case, the right side is classified as “ambiguous”. The full subcategory assigned to the outcome is the pair of classifications assigned to its two break sides. Half-SSTR outcomes analyzed in Figures 7 and S7 consist of outcomes classified as “intended” on the side corresponding to the 3′ end of the donor and “unintended” on the 5′ end of the donor. Capture of donor fragment outcomes consist of outcomes classified as “unintended” on both sides.

If the gap can be covered by a single alignment to the human genome or to the cow genome, the outcome is assigned category “genomic insertion”, with the subcategory indicating which organism the gap covering alignment was from.

If none of the above conditions are satisfied but the outcome architecture contains two target edge alignments with an unexplained gap in between, the outcome is classified as “insertion” if there is no missing reference sequence on the target between the target edge alignments, and as “insertion with deletion” if there is.

If none of these conditions are satisfied, the outcome is assigned to category “uncategorized”. Finally, because many of the individual repair outcome sequences seen in a screen will occur many times each, performing the categorization process described above on each outcome sequence would result in substantial computational redundancy. To improve performance, data from a screen is first preprocessed to identify any sequences that occur more than once across all UMIs for all CRISPRi sgRNAs. The categorization process described above is applied to each common sequence to populate a lookup table of common sequence to outcome. Then, all sequences for each UMIs for each CRISPRi sgRNA are categorized by first checking this lookup table for each sequence and only carrying out the categorization process on sequences not present in the table.

Following categorization of all reads from a screen, the number of occurrences of each combination of outcome category, subcategory, and details is counted for each CRISPRi sgRNA. Outcome counts for the entire screen are then collected into a matrix with rows for each outcome (category, subcategory, and details) combination and columns for each CRISPRi perturbation.

### Statistical significance in volcano plots

Statistical tests consist of asking whether knockdown of a gene significantly changes the fraction of repair outcomes that occupy a given category. To test the null hypothesis that knockdown does not change the fraction of outcomes compared to non-targeting sgRNAs, accounting for the fact that a subset of sgRNAs may achieve weak or no knockdown and therefore that different sgRNAs for the same gene may vary in phenotypic strength, we first calculate a single-sgRNA p-value for each sgRNA targeting a gene, then combine these individual p-values into a gene-level p-value. This combination procedure allows either a single sgRNA with a sufficiently strong phenotype or multiple sgRNAs with weaker but consistent phenotypes to achieve gene-level significance.

To calculate gene-level p-values for a given gene, set of outcomes, and direction of effect (increasing or decreasing), the baseline fraction of relevant outcomes in unperturbed cells was first calculated across all UMIs assigned to non-targeting sgRNAs. Call this quantity f_nt_. Suppose there are g sgRNAs targeting the gene. For each such sgRNA, a single sgRNA p-value was then calculated as the probability that Binomial(d, f_nt_) is less than or equal to n (if the direction being tested is decreasing) or greater than or equal to n (if the direction being tested is increasing), where n is the number of UMIs for the sgRNA with outcomes in the relevant outcome set, and d is the total number of UMIs for the sgRNA. Note that this process may overestimate the statistical significance of a change in outcome frequency if each UMI does not represent a distinct editing outcome. This could happen if multiple daughter cells of the same mother cell (produced by cell division in the time between finishing edit installation and harvesting) are captured. It could also happen if distinct UMIs are artificially assigned to the same editing outcome, either by recombination during sequencing library preparation or by the introduction of a mutation in a UMI sequence at an early enough PCR cycle to produce sufficient reads reporting both the original and mutated UMI sequences.

To combine the g single sgRNA p-values into a gene-level p-value, the p-values were sorted into increasing order. For each k from 1 to g, the probability of observing k independent p-values at least as small as the k’th smallest out of g attempts was calculated as the probability that at least k out of n independently generated uniform (0, 1) deviates are all less than or equal to the rank k single sgRNA p-value. The minimum probability produced by these g tests was retained. This minimum value was multiplied by a Bonferonni correction factor of g to conservatively correct for the fact that this represents testing g (not completely independent) hypotheses.

After carrying out this process for both possible effect directions (increasing and decreasing), the relevant direction of effect for the gene was taken to be the direction with the minimum p-value, and this p-value was converted to a two-sided test by multiplying this minimum p-value by 2.

### Ranking sgRNAs by outcome redistribution activity

To rank CRISPRi sgRNAs in a screen by overall outcome redistribution activity, a chi-squared-like statistic is calculated to quantify the statistical strength of the extent to which editing outcomes are redistributed by each sgRNA relative to all non-targeting sgRNAs. Given a targeting sgRNA t and a set of n relevant outcomes, let n_t, i_ be the number of UMIs for outcome i in sgRNA t and n_nt, i_ by the number of UMIs for outcome i collectively across all non-targeting sgRNAs. Let 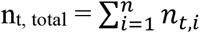 and 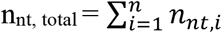. Then the expected number of counts for outcome i in sgRNA t if sgRNA t has no effect is e_t, i_ = n_nt, i_ / n_nt, total_ * n_t, total_. The chi-squared statistic for sgRNA t is then 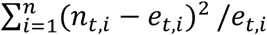.

### Clustering

In hierarchical clustering of data from a single screen or composite data from multiple screens, sgRNAs were ranked by total outcome redistribution activity across all outcomes above indicated baseline frequency thresholds as described above, and the indicated number of most active were retained. Clustering of outcomes was performed using the scipy.cluster.hierarchy.linkage function from scipy 1.5.4 using the ‘correlation’ metric and single linkage with the optional_ordering argument set to True. Clustering of sgRNAs was performed using HDBSCAN version 0.8.26, using pre-computed cosine distances with a minimum cluster size of 2, minimum samples of 1, and cluster selection epsilon of 0.02.

### Dimensionality reduction

Dimensionality reduction of composite Cas9 repair maps was performed using umap-learn version 0.4.6^86^.

### Perturb-seq analysis

Data from the Perturb-seq experiment was analyzed using custom Python scripts built on SCANPY^87^. Assignments of CRISPRi sgRNAs to cells was performed using the mixture model approach described in^57^. Assignment of cells to cell cycle stages was performed as described in the SCANPY documentation at https://nbviewer.jupyter.org/github/theislab/scanpy_usage/blob/master/180209_cell_cycle/cell_cycle.ipynb. Briefly, each cell is scored for expression of a set of S-phase marker genes and a set of G2/M-phase marker genes and assigned to S or G2/M if either of these sets are highly expressed, or to G1 if neither are. Marker gene sets for cell cycle stages were taken from https://github.com/hbc/tinyatlas/blob/master/cell_cycle/Homo_sapiens.csv, which is derived from gene lists originally describe in^88^.

