## SUPPLEMENTARY MATERIALS for "Mapping the Genetic Landscape of DNA Double-strand Break Repair"

### *Supplementary Figures and Legends*

Hussmann *et al* **Figure S1**

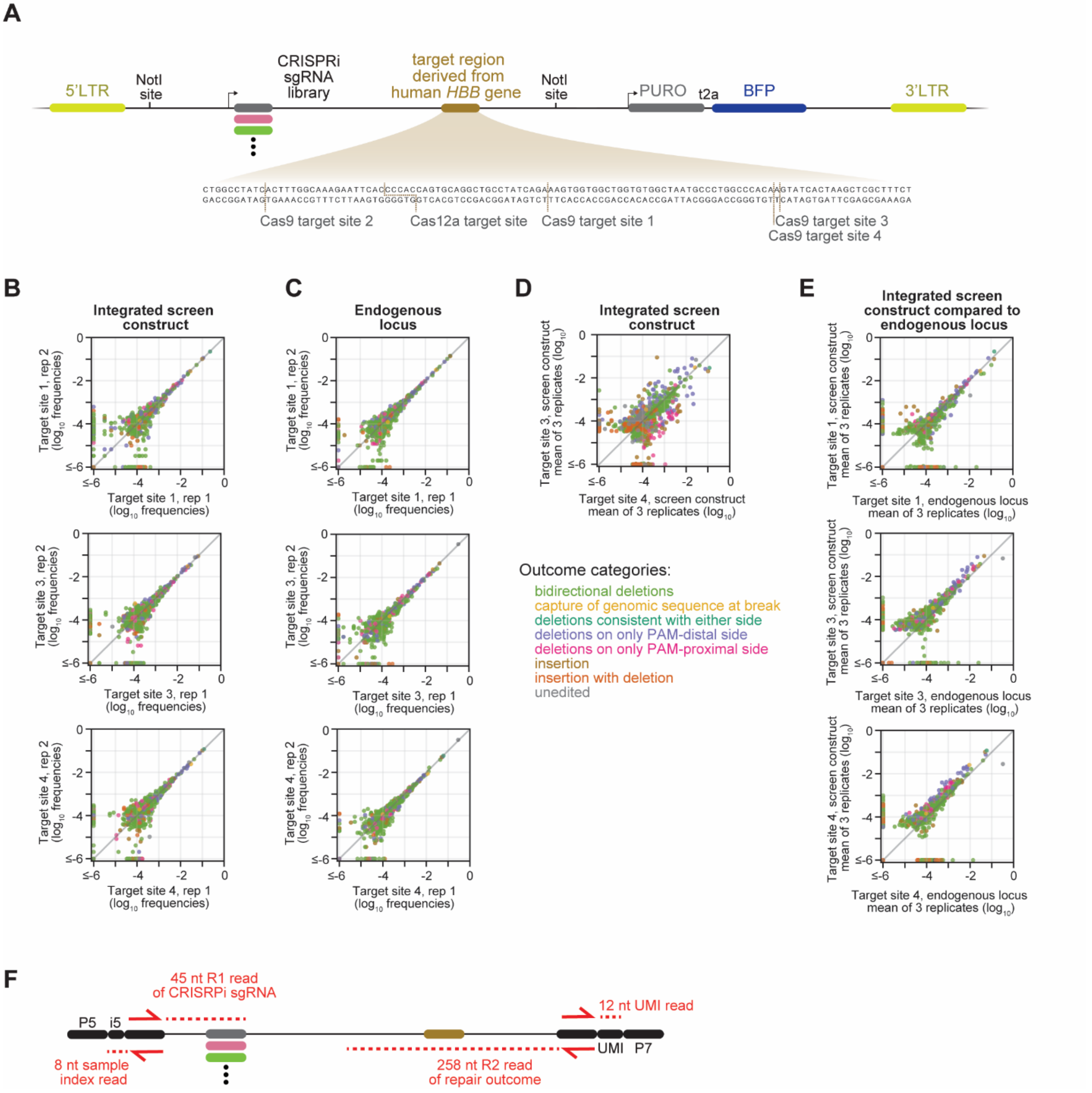

**Figure S1. Design and validation of Repair-seq screen vector and sequencing strategy**

(A) Schematic of lentiviral Repair-seq screen vector. Elements are not represented at scale. LTR indicates long terminal repeat.

(B) Comparisons between replicates (rep) in the frequencies of individual repair outcomes at sites targeted by RNPs (top, Cas9 target site 1; middle, Cas9 target site 3; bottom, Cas9 target site 4; Table S1) within the context of the genomically integrated Repair-seq construct in K562 cells. In each panel, each dot represents a single sequence outcome, with dots colored according to sequence architecture category. Locations of relevant target sites are indicated in panel A.

(C) Comparisons between replicates (rep) in the frequencies of individual repair outcomes at sites targeted by RNPs (top, Cas9 target site 1; middle, Cas9 target site 3; bottom, Cas9 target site 4; Table S1) at the endogenous *HBB* locus. In each panel, each dot represents a single sequence outcome, with dots colored according to sequence architecture category.

(D) Comparison of the frequencies of individual repair outcomes at 2 different Cas9 target sites that are offset from each other by a single nt. Outcomes were measured from DSBs induced in the context of the genomically integrated Repair-seq construct.

(E) Comparison between the frequencies of individual repair outcomes generated within the genomically integrated Repair-seq construct (x-axis, mean of 3 replicates) and the endogenous locus (y-axis, mean of 3 replicates) for each of 3 different Cas9 target sites (top, Cas9 target site 1; middle, Cas9 target site 3; bottom, Cas9 target site 4; Table S1).

(F) Schematic of sequencing strategy. Read 1 (R1, 45 nt) was used to read out the CRISPRi sgRNA sequence, Read 2 (R2, 258 nt) was used to read out the repair outcome. One short index read is used to read the unique molecular identifier (UMI), which is ligated onto molecules before amplification. Another short index read is used to read a sample index (i5), allowing multiplexing of libraries from multiple screens. Elements not represented at scale.

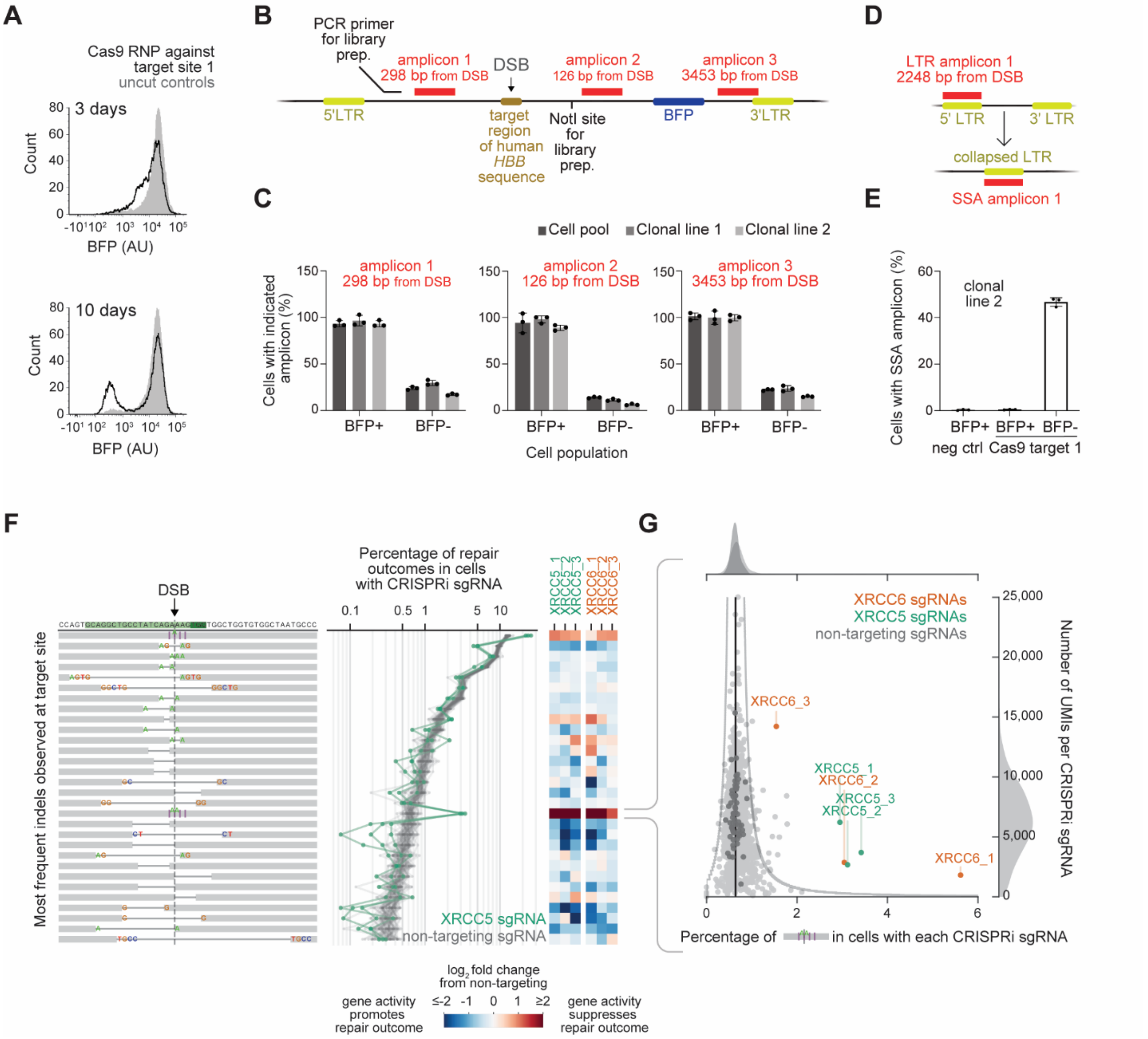
A
Cas9 RNP against target site 1
uncut controls
3 days
10 days
Count
BFP (AU)
B
PCR primer for library prep.
amplicon 1 298 bp from DSB
DSB
amplicon 2 126 bp from DSB
amplicon 3 3453 bp from DSB
5'LTR
target region of human HBB sequence
NotI site for library prep.
BFP
3'LTR
C
Cell pool
Clonal line 1
Clonal line 2
Cells with indicated amplicon (%)
BFP+
BFP-
Cell population
D
LTR amplicon 1 2248 bp from DSB
5' LTR
3' LTR
collapsed LTR
SSA amplicon 1
E
Cells with SSA amplicon (%)
clonal line 2
neg ctrl
Cas9 target 1
F
DSB
Most frequent indels observed at target site
Percentage of repair outcomes in cells with CRISPRi sgRNA
XRCC5_1
XRCC5_2
XRCC5_3
XRCC6_1
XRCC6_2
XRCC6_3
XRCC5 sgRNA
non-targeting sgRNA
log2 fold change from non-targeting
gene activity promotes repair outcome
gene activity suppresses repair outcome
G
XRCC6 sgRNAs
XRCC5 sgRNAs
non-targeting sgRNAs
Number of UMIs per CRISPRi sgRNA
Percentage of in cells with each CRISPRi sgRNA

**Figure S2. Characterization of observed and unobserved repair outcomes**

(A) Distributions of BFP fluorescence in K562 cells expressing dCas9-KRAB and transduced with AC001 library measured 3 days (top) or 10 days (bottom) after electroporation of Cas9 RNP against target site 1.

(B) Schematic depicting the locations of ddPCR amplicons and sequences required for Repair-seq library preparation (PCR primer and NotI site) on the screening construct. Indicated distances are relative to the DSB on the *HBB* target region (site 1). Elements are not represented at scale.

(C) Quantification of the indicated amplicons detected in BFP+ and BFP- cell populations isolated by fluorescence activated cell sorting 4 days after electroporation of an RNP targeting *HBB* site 1. Percentage of cells with each intact amplicon was calculated by comparing the normalized amplicon concentration from indicated sample to that of cells electroporated with gRNA-neg RNP.

(D) Schematic depicting the "LTR amplicon," the inferred single-strand annealing (SSA) repair event, and the resulting "SSA amplicon". Elements are not represented at scale.

(E) Quantification of the "SSA amplicon" detected in BFP+ and BFP- cell populations isolated by fluorescence activated cell sorting 4 days after electroporation of the indicated RNPs. Measurements were normalized to an *ACTB* control amplicon from each sample and compared to the normalized concentration of the "LTR amplicon" in cells electroporated with gRNA-neg RNP (Table S1).

(F) Data from initial Repair-seq screen highlighting phenotypes of *XRCC5* and *XRCC6*, including diagrams (left) of the 30 most frequent indels observed in cells receiving non-targeting CRISPRi sgRNAs, the frequency (middle) of each outcome in the presence of individual non-targeting sgRNAs (60 grey lines) or sgRNAs targeting *XRCC5* (3 green lines), and heatmaps (right) of $\log_2$ fold changes in the frequency of each outcome in the presence of *XRCC5*- and *XRCC6*-targeting sgRNAs relative to the average frequency across all non-targeting sgRNAs.

(G) Effect of every sgRNA in initial Repair-seq screen on the frequency of the indicated 2 nt insertion at the break. For each sgRNA, the number of UMIs recovered for that sgRNA (y-axis) is plotted against the percentage of UMIs reporting this specific deletion (x-axis). Green and orange dots mark sgRNAs targeting *XRCC5* and *XRCC6*, dark grey dots mark individual non-targeting sgRNAs, and light grey dots mark all other sgRNAs. Black vertical line indicates the average percentage across all non-targeting sgRNAs. Dark grey density plot above the scatter plot shows a kernel density estimate (KDE) of the distribution of frequencies in non-targeting sgRNAs and light grey density plots above and to the right show KDEs of the distribution of frequencies (top) and UMI counts (right) for all sgRNAs.

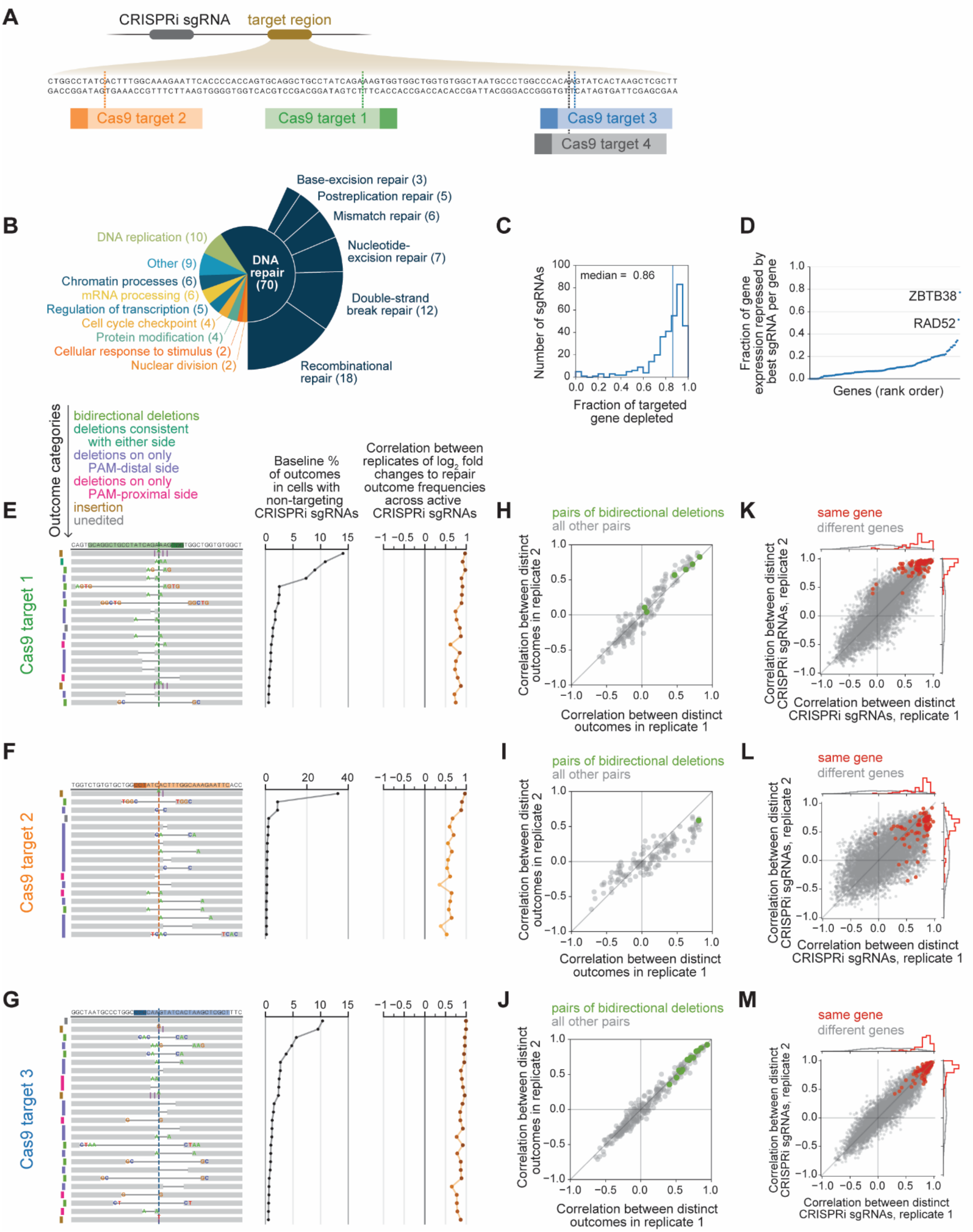
A
CRISPRi sgRNA
target region
Cas9 target 2
Cas9 target 1
Cas9 target 3
Cas9 target 4
B
DNA repair (70)
Base-excision repair (3)
Postreplication repair (5)
Mismatch repair (6)
Nucleotide-excision repair (7)
Double-strand break repair (12)
Recombinational repair (18)
DNA replication (10)
Other (9)
Chromatin processes (6)
mRNA processing (6)
Regulation of transcription (5)
Cell cycle checkpoint (4)
Protein modification (4)
Cellular response to stimulus (2)
Nuclear division (2)
C
median = 0.86
Number of sgRNAs
Fraction of targeted gene depleted
D
Fraction of gene expression repressed by best sgRNA per gene
ZBTB38
RAD52
Genes (rank order)
E
Outcome categories
bidirectional deletions
deletions consistent with either side
deletions on only PAM-distal side
deletions on only PAM-proximal side
insertion
unedited
Baseline % of outcomes in cells with non-targeting CRISPRi sgRNAs
Correlation between replicates of log2 fold changes to repair outcome frequencies across active CRISPRi sgRNAs
Cas9 target 1
F
Cas9 target 2
G
Cas9 target 3
H
I
J
pairs of bidirectional deletions
all other pairs
Correlation between distinct outcomes in replicate 1
Correlation between distinct outcomes in replicate 2
K
L
M
same gene
different genes
Correlation between distinct CRISPRi sgRNAs, replicate 1
Correlation between distinct CRISPRi sgRNAs, replicate 2

**Figure S3. Validation of CRISPRi knockdown by sub-library and of the reproducibility of outcome redistribution profiles**

(A) Relative locations of Cas9 target sites within target region of screen construct.

(B) Functional annotation classes for genes targeted by CRISPRi sub-library (see also Table S6).

(C) Histogram of distribution of fractional knockdowns of targeted genes produced by sgRNAs in CRISPRi sub-library as measured by Perturb-seq. For each CRISPRi sgRNA, a pseudo-bulk expression profile for the sgRNA is assembled by summing UMIs counts from all cells that received only that sgRNA, and the expression of the gene targeted by the sgRNA in this profile is compared to expression of the gene in a profile for cells that received non-targeting sgRNAs.

(D) Distribution of the highest fractional knockdown of each gene achieved across all sgRNAs targeting the gene in CRISPRi sub-library. Genes are ordered by highest fractional knockdown. ZBTB38 and RAD52 are the most extreme outliers for which effective knockdown is not achieved by any sgRNA.

(E+F+G) Diagrams (left) depict the most frequent repair outcomes observed for Cas9 target site 1 (E), target site 2 (F), and target site 3 (G). Graphs show baseline percentages of each outcome across all non-targeting sgRNAs, averaged across two biological replicates (middle), and correlation between replicates in $\log_2$ fold changes across all sgRNAs that produce a significant overall redistribution of outcomes for each depicted outcome (right).

(H+I+J) Correlations between outcome signatures ($\log_2$ fold changes in frequency of an outcome across active sgRNAs) for pairs of distinct outcomes in replicate 1 (x-axis) and replicate 2 (y-axis) for target site 1 (H), target site 2 (I), and target site 3 (J). Green points mark pairs of distinct bidirectional deletions; light grey points mark all other outcome pairs.

(K+L+M) Correlations between sgRNA signatures ($\log_2$ fold changes produced by a CRISPRi sgRNA in the frequency of each outcome present above baseline frequency of 0.5%) for distinct CRISPRi sgRNAs in replicate 1 (x-axis) and replicate 2 (y-axis) screens at Cas9 target site 1 (K), Cas9 target site 2 (L), and Cas9 target site 3 (M). Red points mark pairs of sgRNAs targeting the same gene; grey points mark pairs of sgRNAs targeting distinct genes.

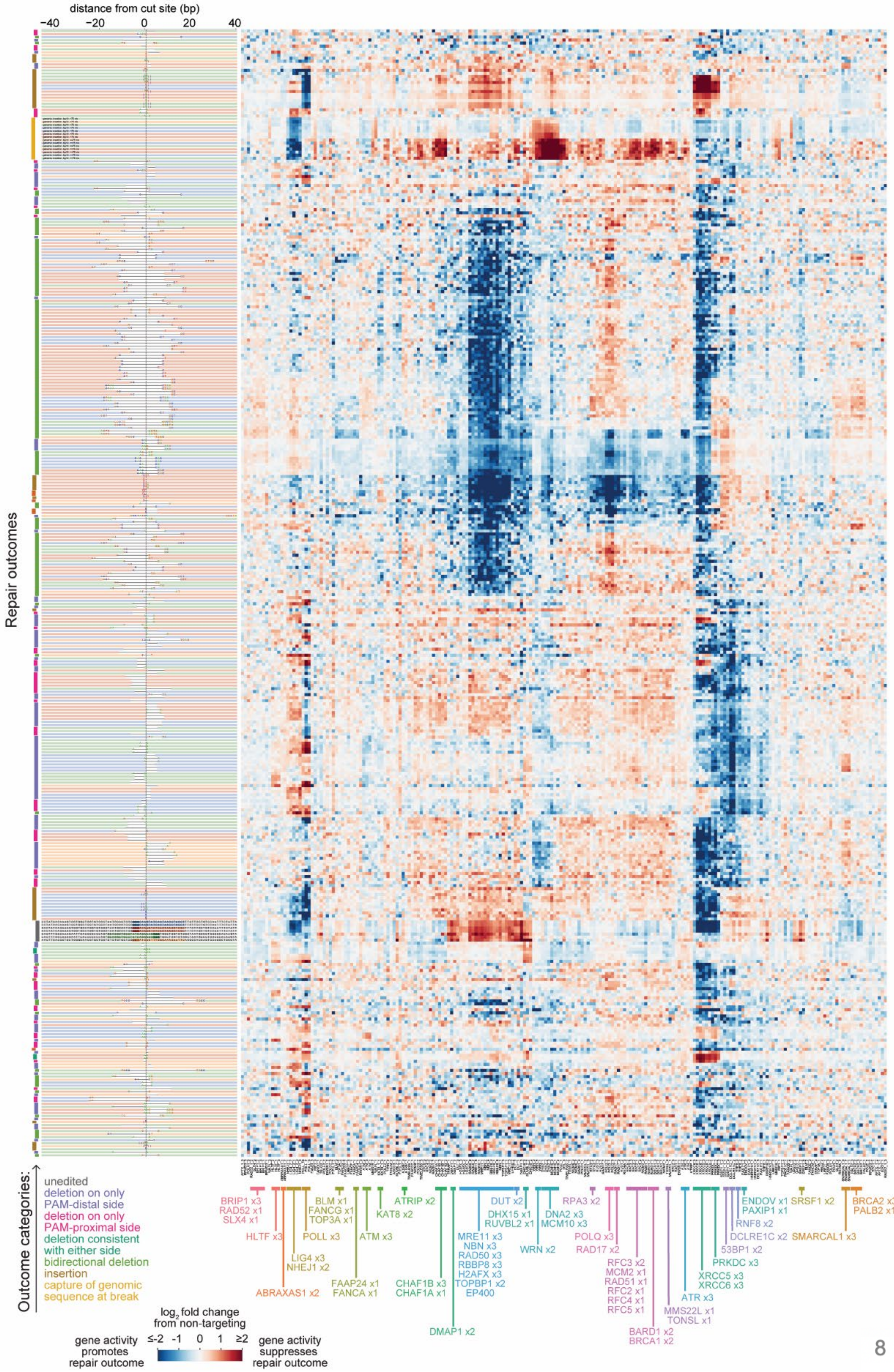
distance from cut site (bp)
−40
−20
0
20
40
Repair outcomes
Outcome categories:
unedited
deletion on only PAM-distal side
deletion on only PAM-proximal side
deletion consistent with either side
bidirectional deletion
insertion
capture of genomic sequence at break
log2 fold change from non-targeting
gene activity promotes repair outcome
≤-2
-1
0
1
≥2
gene activity suppresses repair outcome
BRIP1 x3
RAD52 x1
SLX4 x1
HLTF x3
ABRAXAS1 x2
LIG4 x3
NHEJ1 x2
BLM x1
FANCG x1
TOP3A x1
POLL x3
FAAP24 x1
FANCA x1
ATM x3
KAT8 x2
ATRIP x2
CHAF1B x3
CHAF1A x1
DMAP1 x2
MRE11 x3
NBN x3
RAD50 x3
RBBP8 x3
H2AFX x3
TOPBP1 x2
EP400
DUT x2
DHX15 x1
RUVBL2 x1
DNA2 x3
MCM10 x3
WRN x2
RPA3 x2
POLQ x3
RAD17 x2
RFC3 x2
MCM2 x1
RAD51 x1
RFC2 x1
RFC4 x1
RFC5 x1
BARD1 x2
BRCA1 x2
MMS22L x1
TONSL x1
ATR x3
XRCC5 x3
XRCC6 x3
PRKDC x3
53BP1 x2
DCLRE1C x2
RNF8 x2
ENDOV x1
PAXIP1 x1
SRSF1 x2
SMARCAL1 x3
BRCA2 x3
PALB2 x1

**Figure S4. Hierarchical clustering of sgRNA signatures and outcomes signatures across screens at four Cas9 target sites**

Composite matrix of $\log_2$ fold changes in outcome frequencies produced by the most active CRISPRi sgRNAs in screens at four Cas9 target sites (2 replicates of target sites 1, 3, and 4, and 1 replicate of target site 2). Rows represent outcomes from individual screen replicates, and columns represent individual CRISPRi sgRNAs, with data hierarchically clustered along both dimensions. All individual outcomes present above baseline frequency of 0.2% in each screen are included, as well as two composite outcome categories per screen representing outcomes in which genomic sequence fragments <75 nts or >75 nts are captured at breaks. Any CRISPRi guide that is among the 100 strongest redistributors of overall outcome frequency in any screen is included. Color shading of outcome diagrams indicates target site of origin (green - target site 1; orange - target site 2; blue - target site 3; red - target site 4). Outcome sequence architecture categories are labelled to the left of diagrams. sgRNAs are labeled by cluster assignments produced by HDBSCAN clustering.

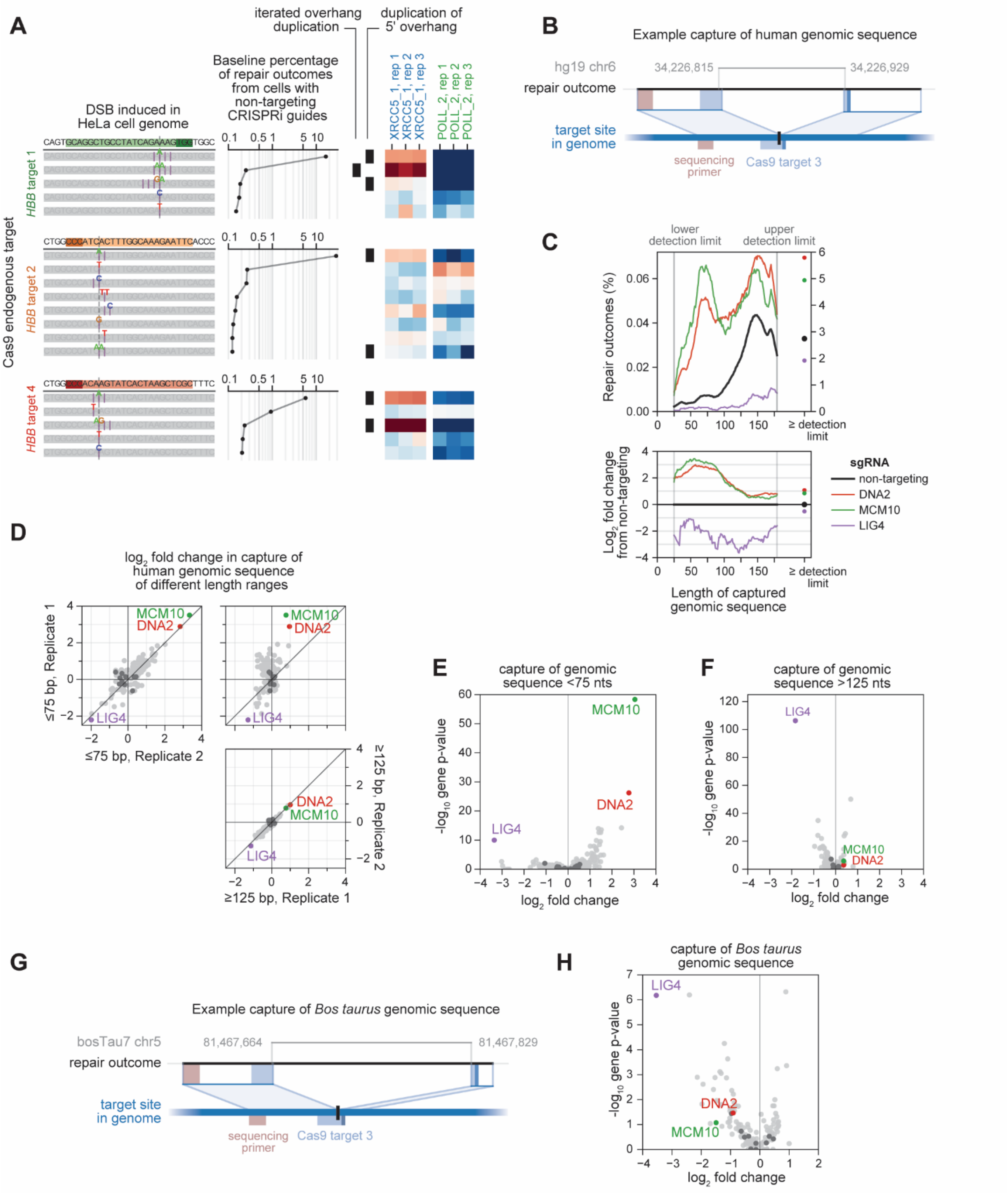
A
DSB induced in HeLa cell genome
Baseline percentage of repair outcomes from cells with non-targeting CRISPRi guides
iterated overhang duplication
duplication of 5' overhang
XRCC5_1, rep 1
XRCC5_1, rep 2
XRCC5_1, rep 3
POLL_2, rep 1
POLL_2, rep 2
POLL_2, rep 3
Cas9 endogenous target
HBB target 1
HBB target 2
HBB target 4
B
Example capture of human genomic sequence
hg19 chr6
34,226,815
34,226,929
repair outcome
target site in genome
sequencing primer
Cas9 target 3
C
lower detection limit
upper detection limit
Repair outcomes (%)
≥ detection limit
Log2 fold change from non-targeting
Length of captured genomic sequence
sgRNA
non-targeting
DNA2
MCM10
LIG4
D
log2 fold change in capture of human genomic sequence of different length ranges
≤75 bp, Replicate 1
≤75 bp, Replicate 2
≥125 bp, Replicate 1
≥125 bp, Replicate 2
E
capture of genomic sequence <75 nts
-log10 gene p-value
log2 fold change
F
capture of genomic sequence >125 nts
G
Example capture of Bos taurus genomic sequence
bosTau7 chr5
81,467,664
81,467,829
H
capture of Bos taurus genomic sequence

**Figure S5. Validation of insertion phenotypes at endogenous loci and characterization of genomic capture outcome**

(A) Effects of repressing *XRCC5* and *POLL* in HeLa cells expressing dCas9-KRAB on insertion frequencies at endogenous *HBB* loci corresponding to Cas9 targets 1, 2, and 4. Diagrams (left) show all insertions observed at baseline frequencies of at least 0.1% in arrayed experiments, with line plots (middle) showing baseline frequencies of each insertion. Heatmaps (right) show log2 fold changes in insertion frequencies produced by one CRISPRi sgRNA targeting each of XRCC5 or POLL in three biological replicates. Annotations to the left of heatmaps mark duplication insertions as in Figure 4F.

(B) Diagram of an example repair outcome in which a segment of human genomic sequence has been inserted at a double-stranded break. Bottom blue line represents the target site in the integrated screen vector, middle black line represents the sequencing read of the repair outcome, blue lines between the target site and the read represent local alignments between the two, and grey line above the read represents a local alignment of the read to chromosome 6 of the human reference genome.

(C) Distribution of lengths of captured human genomic sequence in a screen performed at Cas9 target site 3. This target site is located closer to the sequencing primer, maximizing the range of captured sequence lengths that can be resolved before reaching the end of the sequencing read. Lines in top panel plot the percentage of repair outcomes for which genomic sequence of the indicated length was captured (averaged over 21 nt windows) for all non-targeting sgRNAs (black), all *DNA2* sgRNAs (red), all *MCM10* sgRNAs (green), or all *LIG4* sgRNAs (purple). Lower detection limit indicates the length below which insertion sequence cannot be unambiguously assigned a source, and upper detection limit indicates the length above which the sequencing read is not long enough to resolve the end of an insertion directly at the DSB. The total percentage of outcomes with captured sequence above the upper detection limit is plotted to the far right, with a different y-axis scale. Lines in bottom panel plot $\log_2$ fold changes in length-specific insertion frequency for *DNA2*, *MCM10*, and *LIG4* sgRNAs relative to non-targeting sgRNAs, averaged over 21 nt windows.

(D) Comparison of the effects of gene knockdowns on frequencies of capture of human genomic sequence of different length ranges in screens performed at Cas9 target site 3. To maximize the ability to detect differential genetic modulators of potentially overlapping populations, separated length ranges of less than 5 nts and greater than 125 nts were considered. Gene-level effects were calculated as the mean $\log_2$ fold change in frequency from non-targeting for the two most extreme sgRNAs targeting each gene.

(E+F) Volcano plots of strength and statistical significance of gene-level effects on the frequency of capture of human genomic sequence stretches less than 75 nts (E) or greater than 125 nts (F) in a screen performed at Cas9 target site 1. $\text{Log}_2$ fold changes are the average of the two most extreme sgRNAs targeting each gene, and p-values are calculated with a binomial model for each sgRNA and aggregated into gene-level p-values.

(G) Diagram of an example repair outcome in which a segment of *Bos taurus* genomic sequence has been inserted at a DSB. Bottom blue line represents the target site in the integrated screen vector, middle black line represents the sequencing read of the repair outcome, blue lines between the target site and the read represent local alignments between the two, and grey line above the read represents a local alignment of the read to chromosome 5 of the *Bos taurus* reference genome.

(H) Volcano plot of strength and statistical significance of gene-level effects on the frequency of capture of *Bos taurus* genomic sequence in a screen performed at Cas9 target site 1. $Log_2$ fold changes are the average of the two most extreme sgRNAs targeting each gene, and p-values are calculated with a binomial model for each sgRNA and aggregated into gene-level p-values.

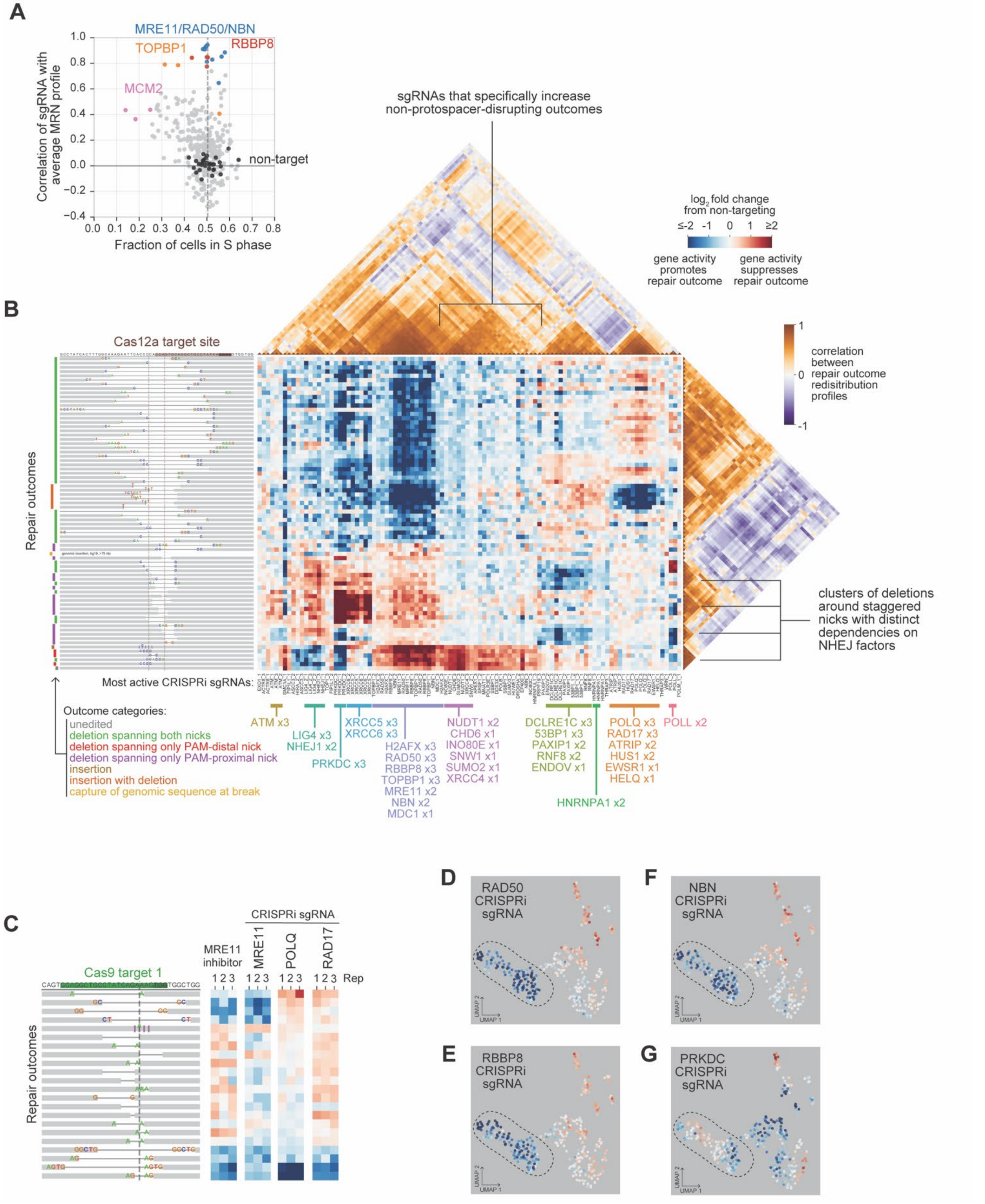
A
MRE11/RAD50/NBN
TOPBP1
RBBP8
MCM2
non-target
Correlation of sgRNA with average MRN profile
Fraction of cells in S phase
B
sgRNAs that specifically increase non-protospacer-disrupting outcomes
log2 fold change from non-targeting
gene activity promotes repair outcome
gene activity suppresses repair outcome
correlation between repair outcome redistribution profiles
Cas12a target site
Repair outcomes
clusters of deletions around staggered nicks with distinct dependencies on NHEJ factors
Most active CRISPRi sgRNAs:
Outcome categories:
unedited
deletion spanning both nicks
deletion spanning only PAM-distal nick
deletion spanning only PAM-proximal nick
insertion
insertion with deletion
capture of genomic sequence at break
ATM x3
LIG4 x3
NHEJ1 x2
PRKDC x3
XRCC5 x3
XRCC6 x3
H2AFX x3
RAD50 x3
RBBP8 x3
TOPBP1 x3
MRE11 x2
NBN x2
MDC1 x1
NUDT1 x2
CHD6 x1
INO80E x1
SNW1 x1
SUMO2 x1
XRCC4 x1
DCLRE1C x3
53BP1 x3
PAXIP1 x2
RNF8 x2
ENDOV x1
HNRNPA1 x2
POLQ x3
RAD17 x3
ATRIP x2
HUS1 x2
EWSR1 x1
HELQ x1
POLL x2
C
Cas9 target 1
Repair outcomes
MRE11 inhibitor
CRISPRi sgRNA
MRE11
POLQ
RAD17
Rep
D
RAD50 CRISPRi sgRNA
F
NBN CRISPRi sgRNA
E
RBBP8 CRISPRi sgRNA
G
PRKDC CRISPRi sgRNA
UMAP 1
UMAP 2

**Figure S6. Data from Cas12a Repair-seq screen and outcome signatures produced by repression of *POLQ* and MRN complex genes**

(A) Comparison of the fraction of all cells containing each CRISPRi sub-library sgRNA assigned to S phase based on transcriptional profiles in Perturb-seq data (x-axis) and the correlation of the same sgRNA's outcome redistribution profile across all Cas9 target sites with the average profile for all sgRNAs targeting *MRE11*, *RAD50*, or *NBN* in Repair-seq data (y-axis). Colored dots mark sgRNAs targeting indicated genes.

(B) Map of repair outcome redistribution profiles in a screen performed using Cas12a to induce DSBs. Central heatmap depicts $\log_2$ fold changes in frequencies of the most frequent repair outcomes (diagrams on left, corresponding to rows) for the 100 most active CRISPRi sgRNAs (columns) relative to the average of all non-targeting CRISPRi sgRNAs, hierarchically clustered along both dimensions. Analyzed outcomes include all individual sequence outcomes present above baseline frequency of 0.2%, as well as one composite outcome representing the combined frequency of all outcomes in which stretches of genomic sequence >75nts were captured at the break. Triangular heatmaps depict correlations between pairs of sgRNAs (above) and between pairs of outcomes (right). sgRNA cluster assignments generated by HDBSCAN are labeled below.

(C) Effects of MRE11 inhibitor (mirin) and CRISPRi-mediated depletion of indicated genes on DSB repair outcomes generated at Cas9 target site 1. Data are from two separate experiments, with mirin data collected in K562 cells and CRISPRi data collected in K562 cells expressing dCas9-KRAB. Heatmaps display $\log_2$ fold changes in outcome frequency for corresponding outcomes. Outcomes are sorted by average $\log_2$ fold change across *POLQ* sgRNAs.

(D+E+F+G) $\text{Log}_2$ fold changes in outcome frequencies produced by indicated CRISPRi sgRNAs overlaid on the Cas9 UMAP outcome embedding.

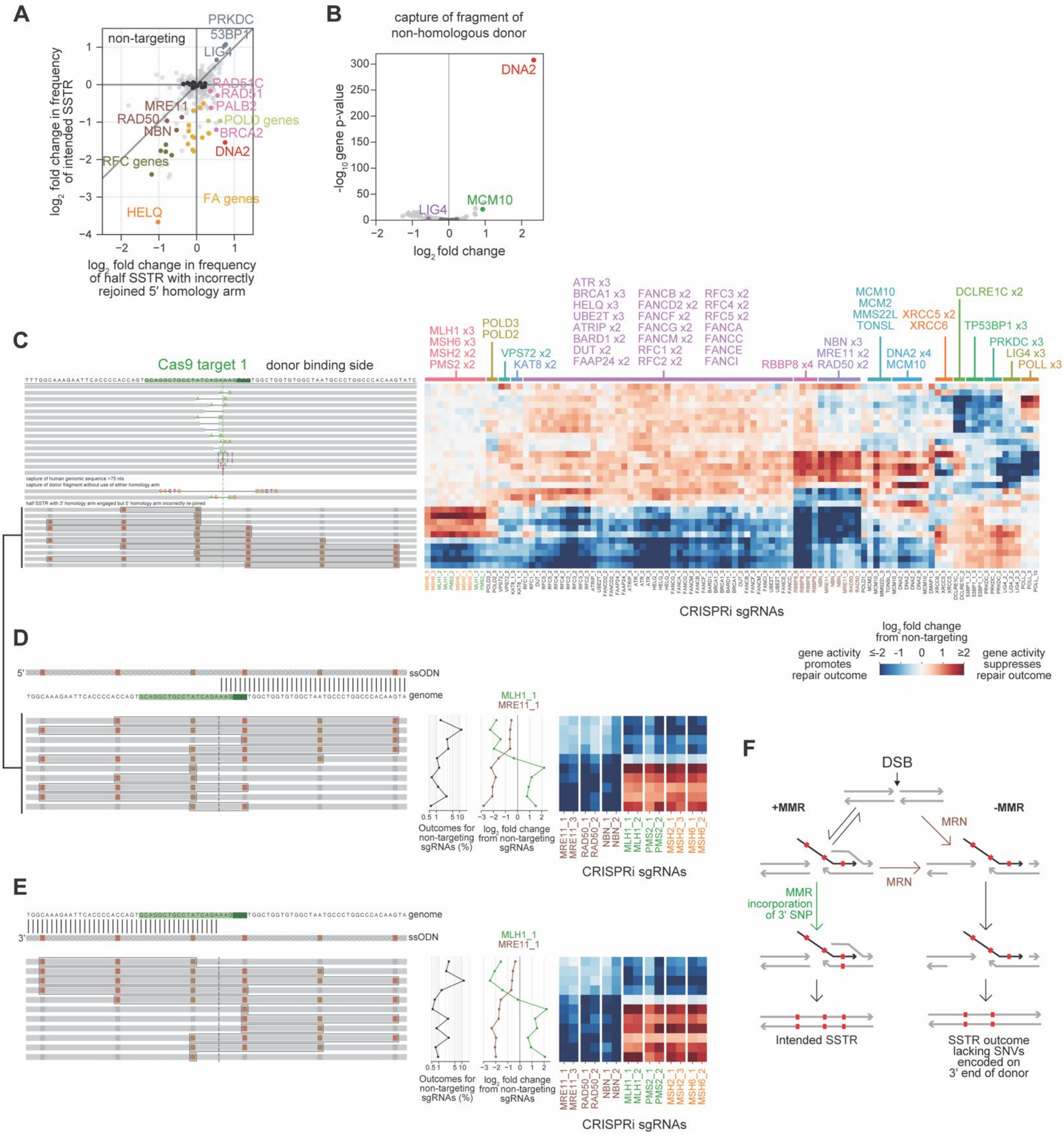
A
non-targeting
PRKDC
53BP1
LIG4
RAD51C
RAD51
PALB2
POLD genes
BRCA2
DNA2
MRE11
RAD50
NBN
RFC genes
FA genes
HELQ
log2 fold change in frequency of intended SSTR
log2 fold change in frequency of half SSTR with incorrectly rejoined 5′ homology arm
B
capture of fragment of non-homologous donor
DNA2
MCM10
LIG4
-log10 gene p-value
log2 fold change
C
Cas9 target 1
donor binding side
capture of human genomic sequence >75 nts
capture of donor fragment without use of either homology arm
half SSTR with 3′ homology arm engaged but 5′ homology arm incorrectly re-joined
MLH1 x3
MSH6 x3
MSH2 x2
PMS2 x2
POLD3
POLD2
VPS72 x2
KAT8 x2
ATR x3
BRCA1 x3
HELQ x3
UBE2T x3
ATRIP x2
BARD1 x2
DUT x2
FAAP24 x2
FANCB x2
FANCD2 x2
FANCF x2
FANCG x2
FANCM x2
RFC1 x2
RFC2 x2
RFC3 x2
RFC4 x2
RFC5 x2
FANCA
FANCC
FANCE
FANCI
RBBP8 x4
NBN x3
MRE11 x2
RAD50 x2
MCM10
MCM2
MMS22L
TONSL
DNA2 x4
MCM10
XRCC5 x2
XRCC6
DCLRE1C x2
TP53BP1 x3
PRKDC x3
LIG4 x3
POLL x3
CRISPRi sgRNAs
D
5′
ssODN
genome
MLH1_1
MRE11_1
Outcomes for non-targeting sgRNAs (%)
log2 fold change from non-targeting sgRNAs
MRE11_1
MRE11_3
RAD50_1
RAD50_2
NBN_1
NBN_2
MLH1_1
MLH1_2
PMS2_1
PMS2_2
MSH2_1
MSH2_3
MSH6_1
MSH6_2
CRISPRi sgRNAs
log2 fold change from non-targeting
gene activity promotes repair outcome
≤-2 -1 0 1 ≥2
gene activity suppresses repair outcome
E
genome
3′
ssODN
MLH1_1
MRE11_1
Outcomes for non-targeting sgRNAs (%)
log2 fold change from non-targeting sgRNAs
CRISPRi sgRNAs
F
DSB
+MMR
-MMR
MRN
MRN
MMR incorporation of 3′ SNP
Intended SSTR
SSTR outcome lacking SNVs encoded on 3′ end of donor

**Figure S7. A diversity of mechanisms incorporate ssODN sequences at DSBs**

(A) Comparison of the effects of gene knockdowns on the combined frequency of all scarless SSTR outcomes (y-axis) and the combined frequency of all half-SSTR outcomes in which the 3′ HA is correctly engaged but the outcome sequence does not rejoin the genome at the intended 5′ HA. Data is from a screen performed using the same donor as in Figure 7B. Gene categories indicated include DNA polymerase δ (POLD) genes, Fanconi anemia (FA) genes, and replication factor C (RFC) genes.

(B) Volcano plot of the effects of gene knockdowns on the combined frequency of capture of donor fragments in a screen performed using an ssODN that has no homology to the target site. Plot depicts $\log_2$ fold change in combined outcome frequency for the two most extreme sgRNAs targeting each gene (x-axis) and an aggregate binomial p-value for all sgRNAs targeting the gene (y-axis). Dark grey dots represent random sets of three individual non-targeting sgRNAs.

(C) Hierarchical clustering of $\log_2$ fold changes produced by the 100 most active CRISPRi sgRNAs in a screen performed with an ssODN containing 6 programmed SNVs evenly spaced around the DSB site. All individual sequence outcomes above 0.5% baseline frequency are included, as well as composite categories representing capture of fragments of human genomic sequence >75 nts, capture of fragments of donor sequence, and half-SSTR outcomes in which the 5′ HA is incorrectly engaged.

(D+E) Effects of knockdown of MRN components or MMR genes on the frequency of scarless SSTR outcomes incorporating different subsets of donor-encoded SNVs. Panels show data from separate screens performed using donors with different strand orientations. Diagrams (top left of each panel) depict the integrated target site along with the strand orientation and programmed edit patterns of the donor. Below, all scarless SSTR outcomes observed at a baseline frequency above 0.5% in each screen are shown. Black line plots show baseline frequencies of each outcome. Brown and green line plots and heatmaps show $\log_2$ fold changes in outcome frequency produced by indicated CRISPRi sgRNAs. Screens were performed at Cas9 target site 1 with an ssODN encoding 6 SNVs oriented such that it can anneal to the PAM-proximal side of a resected DSB (D) or at Cas9 target site 1 with an ssODN that has the reverse complement sequence as in D and therefore can anneal to the PAM-distal side of a resected DSB (E).

(F) Model for roles of MRN complex and mismatch repair (MMR) in mediating scarless SSTR. Partially-complementary ssODNs can anneal to only one side of a resected break, forming heteroduplexes at locations of programmed edits. Consistent with previous models, Repair-seq phenotypes suggest that these heteroduplexes can be processed by MMR, resulting in the installation of SNVs from the annealed side of the donor into the genome and concomitant suppression of scarless SSTR outcomes lacking these SNV. Phenotypes of genes encoding the MRN complex (*MRE11*, *NBN*, *RAD50*) suggest an extension to this model wherein suppressed SSTR outcomes have a strong requirement for resection.

***Supplementary Tables.***

**Table S1.** gRNAs used for DSB-induction.
**Table S2.** CRISPRi library with 1,573 sgRNAs targeting 476 genes.
**Table S3.** Repair-seq screen details.
**Table S4.** CRISPRi library with 366 sgRNAs targeting 118 genes.
**Table S5.** Functional annotation of genes targeted by 1,573 sgRNA library.
**Table S6.** Functional annotation of genes targeted by 366 sgRNA library.
**Table S7.** Oligonucleotides and TaqMan probes used in this study. Within this table * represents a phosphorothioated DNA base.

***Supplementary Sequences.***

**SpCas9.**MKHHHHHHMPKKKRKVMDKKYSIGLDIGTNSVGWAVITDEYKVPSKKFKVLGNTDR
HSIKKNLIGALLFDSGETAEATRLKRTARRRYTRRKNRICYLQEIFSNEMAKVDDSFFHRLEESF
LVEEDKKHERHPIFGNIVDEVAYHEKYPTIYHLRKKLVDSTDKADLRLIYLALAHMIKFRGHFLI
EGDLNPDNSDVDKLFIQLVQTYNQLFEENPINASGVDAKAILSARLSKSRRLENLIAQLPGEKKN
GLFGNLIALSLGLTPNFKSNFDLAEDAKLQLSKDTYDDDLDNLLAQIGDQYADLFLAAKNLSD
AILLSDILRVNTEITKAPLSASMIKRYDEHHQDLTLLKALVRQQLPEKYKEIFFDQSKNGYAGYI
DGGASQEEFYKFIKPILEKMDGTEELLVKLNREDLLRKQRTFDNGSIPHQIHLGELHAILRRQED
FYPFLKDNREKIEKILTFRIPYYVGPLARGNSRFAWMTRKSEETITPWNFEEVVDKGASAQSFIE
RMTNFDKNLPNEKVLPKHSLLYEYFTVYNELTKVKYVTEGMRKPAFLSGEQKKAIVDLLFKTN
RKVTVKQLKEDYFKKIECFDSVEISGVEDRFNASLGTYHDLLKIIKDKDFLDNEENEDILEDIVLT
LTLFEDREMIEERLKTYAHLFDDKVMKQLKRRRYTGWGRLSRKLINGIRDKQSGKTILDFLKSD
GFANRNFMQLIHDDSLTFKEDIQKAQVSGQGDSLHEHIANLAGSPAIKKGILQTVKVVDELVKV
MGRHKPENIVIEMARENQTTQKGQKNSRERMKRIEEGIKELGSQILKEHPVENTQLQNEKLYLY
YLQNGRDMYVDQELDINRLSDYDVDHIVPQSFLKDDSIDNKVLTRSDKNRGKSDNVPSEEVVK
KMKNYWRQLLNAKLITQRKFDNLTKAERGGLSELDKAGFIKRQLVETRQITKHVAQILDSRMN
TKYDENDKLIREVKVITLKSKLVSDFRKDFQFYKVREINNYHHAHDAYLNAVVGTALIKKYPK
LESEFVYGDYKVYDVRKMIAKSEQEIGKATAKYFFYSNIMNFFKTEITLANGEIRKRPLIETNGE
TGEIVWDKGRDFATVRKVLSMPQVNIVKKTEVQTGGFSKESILPKRNSDKLIARKKDWDPKKY
GGFDSPTVAYSVLVVAKVEKGKSKKLKSVKELLGITIMERSSFEKNPIDFLEAKGYKEVKKDLII
KLPKYSLFELENGRKRMLASAGELQKGNELALPSKYVNFLYLASHYEKLKGSPEDNEQKQLFV
EQHKHYLDEIIEQISEFSKRVILADANLDKVLSAYNKHRDKPIREQAENIIHLFTLTNLGAPAAFK
YFDTTIDRKRYTSTKEVLDATLIHQSITGLYETRIDLSQLGGD

**His-AsCpf1-**
**2xNLS_H800A.**MGHHHHHHGSTQFEGFTNLYQVSKTLRFELIPQGKTLKHIQEQGFIEEDKARND
HYKELKPIIDRIYKTYADQCLQLVQLDWENLSAAIDSYRKEKTEETRNALIEEQATYRNAIHDYF
IGRTDNLTDAINKRHAEIYKGLFKAELFNGKVLKQLGTVTTTEHENALLRSFDKFTTYFSGFYE
NRKNVFSAEDISTAIPHRIVQDNFPKFKENCHIFTRLITAVPSLREHFENVKKAIGIFVSTSIEEVFS
FPFYNQLLTQTQIDLYNQLLGGISREAGTEKIKGLNEVLNLAIQKNDETAHIIASLPHRFIPLFKQI
LSDRNTLSFILEEFKSDEEVIQSFCKYKTLLRNENVLETAEALFNELNSIDLTHIFISHKKLETISSA
LCDHWDTLRNALYERRISELTGKITKSAKEKVQRSLKHEDINLQEIISAAGKELSEAFKQKTSEIL
SHAHAALDQPLPTTLKKQEEKEILKSQLDSLLGLYHLLDWFAVDESNEVDPEFSARLTGIKLEM

EPSLSFYNKARNYATKKPYSVEKFKLNFQMPTLASGWDVNKEKNNGAILFVKNGLYYLGIMP
KQKGRYKALSFEPTEKTSEGFDKMYYDYFPDAAKMIPKCSTQLKAVTAHFQTHTTPILLSNNFI
EPLEITKEIYDLNNPEKEPKKFQTAYAKKTGDQKGYREALCKWIDFTRDFLSKYTKTTSIDLSSL
RPSSQYKDLGEYYAELNPLLYHISFQRIAEKEIMDAVETGKLYLFQIYNKDFAKGHHGKPNLHT
LYWTGLFSPENLAKTSIKLNGQAELFYRPKSRMKRMAARLGEKMLNKKLKDQKTPIPDTLYQE
LYDYVNHRLSHDLSDEARALLPNVITKEVSHEIIKDRRFTSDKFFFHVPITLNYQAANSPSKFNQ
RVNAYLKEHPETPIIGIDRGERNLIYITVIDSTGKILEQRSLNTIQQFDYQKKLDNREKERVAARQ
AWSVVGTIKDLKQGYLSQVIHEIVDLMIHYQAVVVLENLNFGFKSKRTGIAEKAVYQQFEKML
IDKLNCLVLKDYPAEKVGGVLNPYQLTDQFTSFAKMGTQSGFLFYVPAPYTSKIDPLTGFVDPF
VWKTIKNHESRKHFLEGFDFLHYDVKTGDFILHFKMNRNLSFQRGLPGFMPAWDIVFEKNETQ
FDAKGTPFIAGKRIVPVIENHRFTGRYRDLYPANELIALLEEKGIVFRDGSNILPKLLENDDSHAI
DTMVALIRSVLQMRNSNAATGEDYINSPVRDLNGVCFDSRFQNPEWPMDADANGAYHIALKG
QLLLNHLKESKDLKLQNGISNQDWLAYIQELRNGSPKKKRKVGSPKKKRKV
